# Canopy spectral reflectance detects oak wilt at the landscape scale using phylogenetic discrimination

**DOI:** 10.1101/2021.01.17.427016

**Authors:** Gerard Sapes, Cathleen Lapadat, Anna K. Schweiger, Jennifer Juzwik, Rebecca Montgomery, Hamed Gholizadeh, Philip A. Townsend, John A. Gamon, Jeannine Cavender-Bares

## Abstract

The oak wilt disease caused by the invasive fungal pathogen *Bretziella fagacearum* is one of the greatest threats to oak-dominated forests across the Eastern United States. Accurate detection and monitoring over large areas are necessary for management activities to effectively mitigate and prevent the spread of oak wilt. Canopy spectral reflectance contains both phylogenetic and physiological information across the visible near-infrared (VNIR) and short-wave infrared (SWIR) ranges that can be used to identify diseased red oaks. We develop partial least square discriminant analysis (PLS-DA) models using airborne hyperspectral reflectance to detect diseased canopies and assess the importance of VNIR, SWIR, phylogeny, and physiology for oak wilt detection. We achieve high accuracy through a three-step phylogenetic process in which we first distinguish oaks from other species (90% accuracy), then red oaks from white oaks (*Quercus macrocarpa*) (93% accuracy), and, lastly, infected from non-infected trees (80% accuracy). Including SWIR wavelengths increased model accuracy by ca. 20% relative to models based on VIS-NIR wavelengths alone; using a phylogenetic approach also increased model accuracy by ca. 20% over a single-step classification. SWIR wavelengths include spectral information important in differentiating red oaks from other species and in distinguishing diseased red oaks from healthy red oaks. We determined the most important wavelengths to identify oak species, red oaks, and diseased red oaks. We also demonstrated that several multispectral indices associated with physiological decline can detect differences between healthy and diseased trees. The wavelengths in these indices also tended to be among the most important wavelengths for disease detection within PLS-DA models, indicating a convergence of the methods. Indices were most significant for detecting oak wilt during late August, especially those associated with canopy photosynthetic activity and water status. Our study suggests that coupling phylogenetics, physiology, and canopy spectral reflectance provides an interdisciplinary and comprehensive approach that enables detection of forest diseases at large scales. These results have potential for direct application by forest managers for detection to initiate actions to mitigate the disease and prevent pathogen spread.

## 1. Introduction

Invasive tree pathogens are a major threat to forest diversity and function (Evans et al., 2010; Hulcr and Dunn, 2011). The damage caused by invasive species can have negative consequences for ecosystem processes and services, including air and water quality maintenance, nutrient and carbon cycling, wood and food provision, and climate regulation (Cavender-Bares et al., 2019; Díaz et al., 2019; Waller et al., 2020). In North American forests, invasive pathogens and pests that infect trees have had devastating impacts over the last century due to multiple factors, including global trade and climate change (Bergot et al., 2004; Liebhold et al., 1995; Sturrock et al., 2011), leading to the loss or potential loss of multiple foundational canopy species such as American chestnut (*Castanea dentata*), elm and ash species (*Ulmus* and *Fraxinus* spp.), and eastern hemlock (*Tsuga canadiensis*). Protecting ecosystems from the threats of invasive species resulting from globalization and a changing climate is one of the most pressing challenges of our times (Díaz et al., 2019; Liebhold et al., 1995; Waller et al., 2020).

The oak genus (*Quercus*) is under threat from multiple pathogens and is of critical management interest due to its dominance in temperate forests of the Eastern US (Johnson et al., 2019). Oaks rank among the most diverse and important tree lineages in the United States, with 91 oak species comprising nearly 30% of biomass in US temperate forests (Cavender-Bares, 2019). Among the pathogens affecting oaks, oak wilt caused by the fungus *Bretziella fagacearum* (de Beer et al., 2017) is considered one of the most destructive threat to oaks (Appel, 1995; Haight et al., 2011; Wilson and Lindsey, 2005). The oak wilt fungus spreads below-ground from diseased trees to neighboring oaks through networks of grafted roots, thus forming centers (i.e., pockets or foci) of diseased oaks. The pathogen is also transmitted above-ground by nitidulid beetles (family *Nitidulidae*) and oak bark beetles (*Pityophthorus spp*) (Gibbs and French 1980). Multiple species of nitidulid beetles are attracted to spore-producing fungal mats that form on branches and main stems of recently wilted red oaks (Gibbs and French, 1980; Juzwik et al., 2011; Juzwik and French, 1983). On a land parcel to larger scale, oak wilt can be most effectively controlled when oak wilt centers are detected and appropriately treated (Juzwik et al., 2011; Koch et al., 2010). This prevents spread or minimizes disease intensification within a stand or the surrounding landscape. Surveys of large, forested areas to identify suspect diseased trees are time-intensive and require expert training. Such surveys are needed for landscape level oak wilt management efforts. Current operational surveillance of forest landscapes in the Upper Midwest USA utilize aerial surveys conducted with fixed wing aircraft, helicopters, and UAVs (Juzwik, 2009). Other airborne imaging spectrometry offers potential for accurate detection of oak wilt at landscape scales.

Canopy spectral reflectance can potentially be used to detect the physiological decline resulting from oak wilt fungus infection, and thus provide forest managers with a powerful tool. Airborne spectral reflectance and indices derived from reflectance spectra have successfully been used to detect other diseases and insect damage, such as Rapid Ohia Death, Emerald Ash Borer, bark beetles, olive decline due to *Xylella fastidiosa*, and both drought- and *Phytophtora*-induced holm oak decline (Asner et al., 2018; Hornero et al., 2021; Lausch et al., 2013; Ogaya et al., 2015; Pontius et al., 2008, 2005; Zarco-Tejada et al., 2018). To date, spectral indices for oak wilt detection have only been developed for oak seedlings (Fallon et al., 2020). Oaks respond to oak wilt infection by forming balloon-like structures called tyloses that occlude vessels within the xylem (Juzwik and Appel, 2016; Yadeta and Thomma, Bart, 2013). Vessel occlusion potentially blocks or slows the spread of the pathogen but also reduces water transport and limits canopy physiological performance by reducing transpiration and photosynthesis and potentially causing photoinhibition (Fallon et al., 2020; Juzwik and Appel, 2016; Struckmeyer et al., 1954). In red oak species, the fungus is rapidly spread internally in the transpiration stream through large diameter springwood vessels before tylose formation limits the pathogen’s spread. However, the tyloses formed contribute to the development of wilt symptoms. Blockage of vascular conduits by tyloses and metabolites produced by the fungus can lead to declines in transpiration and canopy water content as water supply to the canopy is significantly impaired. Changes in photosynthetic activity, foliar pigment pool sizes, and water status can be detected from canopy spectra (Hanavan et al., 2015; Pontius et al., 2005; Serbin et al., 2015). Changes in foliar pigments, particularly carotenoids, chlorophyll, and those involved in the xanthophyll cycle, have recently been shown to be early markers of holm oak decline (Encinas-Valero et al., 2021). Fallon et al. (2020) identified several spectral wavelengths predictive of oak wilt in greenhouse seedlings that were related to leaf photosynthesis and water status. However, spectral properties of seedlings grown and measured under greenhouse conditions may differ significantly from adult trees grown under natural conditions due to growing conditions (e.g., sun, shade, humidity, and selective filtering of solar radiation by glasshouse materials), ontogeny, canopy position, degree of canopy emergence and other factors (Cavender-Bares et al., 2020; Fernandes et al., 2020; Ollinger, 2011). Hence, it is important to explicitly test the extent to which we can detect oak wilt in natural populations of adult trees using spectral reflectance. Identification of wavelengths associated with physiological function may enable detection of isolated trees with oak wilt that would otherwise remain undetected until oak wilt damage is more extensive.

Oak lineages vary in susceptibility to oak wilt. Consequently, identifying vulnerable oak subgenera is crucial to detect the disease and prevent its spread. White oaks (*Quercus* section *Quercus*), such as *Q. alba* and *Q. macrocarpa*, have narrower vessels (Cavender-Bares and Holbrook, 2001) and may produce tyloses efficiently (Cochard and Tyree, 1990) and in a more targeted manner in response to fungal infection (cf. Yadeta and Thomma, Bart, 2013). This slows the spread or compartmentalizes (cf. Shigo, 1984) the pathogen in infected white oak species (Jacobi and MacDonald, 1980; Koch et al., 2010; Schoenweiss, 1959). Thus, symptoms of oak wilt in white oaks appear as scattered wilt or as dieback in the crown that develops over several to many years. In contrast, red oaks (*Quercus* section *Lobatae*), such as *Q. ellipsoidalis* and *Q. rubra*, have larger diameter springwood vessels and tend to delay tylose formation in response to fungal infection, limiting their effectiveness in halting the spread of the fungus through the vascular system (Juzwik and Appel, 2016; Struckmeyer et al., 1954; Yadeta and Thomma, Bart, 2013). Thus, crown wilt symptoms in red oaks progress rapidly and lead to tree death within the same season or early in the subsequent growing season. The comparatively rapid mortality of red oaks, the common occurrence of intra-specific root grafts, and their common production of spore mats on recently wilted trees contribute to the importance of the red oak lineage in driving disease epidemics in the landscape (Menges and Loucks, 1984). Distinguishing red oaks from white oaks and other species across the landscape is therefore a critical step towards large-scale management of oak wilt. Leaf level and canopy-level modeling approaches using spectroscopic data have previously been successful in distinguishing these lineages in experimental systems and manipulated forest communities (Cavender-Bares et al., 2016; Fallon et al., 2020; Williams et al., 2020). We thus anticipate that it is possible to detect red oaks across the landscape remotely by mapping lineage identities from classification algorithms using airborne spectroscopic imagery. Here, we outline a stepwise phylogenetic approach to remote sensing of oak wilt that entails: 1) identifying trees belonging to the oak genus, 2) identifying oaks belonging to the red oak section, and 3) identifying red oaks infected with oak wilt. The phylogenetic approach consists of three sequential steps: first, pixels are classified as either “oak” or “other species” using an algorithm specifically trained to discriminate oaks and filter out pixels classified as other species. Second, pixels classified as “oak” are classified as either “red oak” or “white oak” using an algorithm specifically trained to discriminate between red oaks and white oaks. After filtering out pixels classified as white oaks, pixels classified as “red oak” are classified as either “diseased” or “healthy red oak” using an algorithm specifically trained to distinguish diseased red oaks from healthy read oaks.

The goal of this study is to identify the optimal spectral range for detection of oak wilt in red oak species (*Q. ellipsoidalis* and *Q. rubra*) across landscapes, find key wavelengths indicative of oak wilt and its host species, and identify common spectral indices that distinguish diseased from healthy read oaks based on physiological processes. Specifically, we compare both full-range (visible, near-infrared, shortwave infrared, VSWIR, 400-2500 nm) and VNIR (visible, near-infrared, 400-1000 nm) imaging spectroscopy and assess their accuracy of oak wilt detection. While the VNIR is sensitive to photosynthetic activity and pigments (Curran et al., 1995; Gamon and Surfus, 1999; Ustin et al., 2009), use of the SWIR provides structural and phenotypic information (Townsend et al., 2013) that is strongly coupled with phylogenetic information (Meireles et al., 2020a) including mesophyll integrity, chemical composition, and canopy water content (Jacquemoud and Ustin, 2001; Ramirez et al., 2015; Romero et al., 2012; Sims and Gamon, 2003). Then, we test the efficacy of spectral vegetation indices known to be sensitive to physiological decline and disease response for their ability to differentiate healthy and diseased trees (Pontius, 2014; Pontius 2020) (Table S1). Spectral indices can increase flexibility in the detection approach because they use only a handful of wavelengths and can be easily calculated across platforms as long as the same wavelengths are present (Pontius, 2014). Spectroscopic models that require hundreds of wavelengths can have limited applicability across platforms when sensor measurement characteristics vary (Castaldi et al., 2018; Crucil et al., 2019; Nouri et al., 2017).

Here, we develop statistical models for oak wilt detection at the landscape scale using airborne spectroscopic imagery collected by two airborne systems (AISA Eagle and AVIRIS-NG) (Gholizadeh et al., 2019; Hamlin et al., 2010) covering different ranges of wavelengths (VNIR and VSWIR, respectively). We coupled on-ground tree identification and status surveys with airborne imaging spectroscopy data to assess the capacity of airborne spectroscopy to detect oak wilt in a temperate, mixed hardwood forest that included adult red oak populations. In doing so, we tested the following hypotheses:

i. Canopy reflectance from airborne spectroscopic imagery can accurately detect oak wilt infected trees in a natural forest landscape;
ii. Detecting red oaks susceptible to oak wilt prior to diseased trees based on spectral features specific to their phylogenetic lineage increases oak wilt detection by removing species outside the oak genus and red oak lineage;
iii. A broad spectral range (VNIR+SWIR) exhibits greater detection accuracy than a narrower spectral range (VNIR only) due to additional spectral information related to phylogenetic identity and plant structure; and
iv. Spectral indices including wavelengths associated with photosynthetic activity, carotenoid, chlorophyll, and xanthophyll pigment content, and canopy water status differentiate diseased red oaks from healthy red oaks.

## 2. Methods

### 2.1 Study area

The study area was the University of Minnesota Cedar Creek Ecosystem Science Reserve (CCESR) (N 45°40’21”, W 93°19’94”). Located in central Minnesota at approx. 280 m above sea level, CCESR has a continental climate with cold winters (January mean -10 °C), hot summers (July mean 22.2 °C), and a mean annual precipitation of 660 mm, spread fairly evenly throughout the year. The vegetation is comprised of a mosaic of uplands dominated by oak savanna, prairie, mixed hardwood forest, and abandoned agricultural fields, with lowlands comprised of ash and cedar swamps, acid bogs, marshes, and sedge meadows. Oak savanna used to be one of the dominant vegetation types in the Cedar Creek area before the European settlement (Grigal et al., 1974). The savannas mainly include two species within the red oak group (*Quercus* section Lobatae, (Denk et al., 2017)), *Quercus ellipsoidalis* and *Q. rubra* (northern pin oak and northern red oak, respectively) and a bur oak, *Q. macrocarpa* (within the white oak group, *Quercus* section Quercus). These two oak groups have different sensitivities to oak wilt, with red oaks being more susceptible, dying within one or two years of infection. The presence of oak wilt fungus has been documented in central Minnesota since the 1940’s where it has led to widespread mortality in forests not treated for the disease. At CCESR, the fungal pathogen oak wilt has spread rapidly in the last decade, leading to exponential increases in the number of standing dead trees as a result of recent mortality (Pellegrini et al., 2021). The diversity of tree species and the widespread presence of active oak wilt make CCESR well suited to assess the capacity of airborne spectroscopy to detect oak wilt in red oaks.

### 2.2 Airborne data collection and tree survey

We collected two airborne imaging spectroscopy datasets across the whole study area on two dates in 2016. The first dataset was collected on 07/22/2016 between 9:08 am and 10:24 am local time using “CHAMP” (the CALMIT Hyperspectral Airborne Monitoring Platform), the University of Nebraska – Lincoln’s (UNL) aircraft operated by UNL’s Center for Advanced Land Management Information Technologies (CALMIT) and equipped with a pushbroom imaging spectrometer (AISA Eagle, Specim, Oulu, Finland). Data were collected at an average flight altitude of 1150 m above ground level in the northwest-southeast direction, yielding a spatial resolution of 0.75 m. The AISA Eagle comprises 488 spectral channels covering 400-982 nm with a spectral resolution of 1.25 nm and a field of view of 37.7° under nadir viewing conditions. To increase the signal-to-noise-ratio of the data, spectral on-chip binning was applied. The final product had 63 bands at ca. 9 nm intervals. The AISA Eagle images were geometrically corrected using aircraft GPS and IMU data in Specim’s CaliGeoPRO software. Radiance data were converted to reflectance using the empirical line correction (Conel et al., 1987) on reflectance measurements collected from three calibration tarps (white, grey and black, with approx. 5%, 10%, and 40% reflectance, respectively; Odyssey, Ennis Fabrics, Edmonton, Alberta, Canada) with a portable spectroradiometer (SVC HR-1024i, Spectra Vista Corporation, Poughkeepsie, NY, USA; 350 – 2500 nm) simultaneous to the overflights. SVC reflectance data were resampled to match the wavelength of airborne data and then used in the empirical line correction approach. The second dataset was collected using the Airborne Visible/Infrared Imaging Spectrometer - Next Generation (AVIRIS-NG) by the National Aeronautics and Space Administration (NASA) on 08/22/2016 starting at 03:43 PM local time at an average flight altitude of 1210 m above ground level in the near West-East direction, yielding a spatial resolution of 0.9 m. AVIRIS-NG comprises 432 spectral channels covering 380-2510 nm with a spectral resolution of 5 nm and a field of view of 36° under nadir viewing conditions. We measured the three calibration tarps with our portable spectroradiometer (SVC HR-1024i, Spectra Vista Corporation, Poughkeepsie, NY, USA; 350 – 2500 nm) during the overflights for empirical line correction. AVIRIS-NG images that were delivered by the NASA Jet Propulsion Laboratory (JPL) were orthorectified and atmospherically corrected to obtain apparent surface reflectance using a radiative transfer approach following Thompson et al. (2015), while the AISA Eagle images were corrected using empirical line correction. Because the aircraft images were acquired from different platforms on different dates, with different instruments and atmospheric correction approaches, our objective was to test the relative capacities of each system rather than to integrate the results from each. Specifically, absolute values from the two sensors are not directly comparable. However, if images from each sensor are processed consistently (see Wang et al., 2021) the results from each set of analyses to the different datasets can be compared. While using two different sensors simultaneously is not necessary to detect oak wilt, not all forest managers have access to all sensor types. Testing two different sensors allows us to find wavelengths and indices predictive of oak wilt for multiple sensors and provides alternative tools for oak wilt management.

About one year after collecting airborne data, between June-August of 2017, we tagged 423 mature trees of 12 species with no visual symptoms of oak wilt in woodland and savanna areas (see Table S2 for a description of the range of heights and diameters at breast height, and the number of trees included for each species) including 47 *Quercus ellipsoidalis* E.J. Hill (red oak section, particularly vulnerable to oak wilt). In addition to the 423 healthy trees, we tagged 41 adult *Q. ellipsoidalis* trees (total of 464 trees) with foliar symptoms characteristic of oak wilt (e.g., leaf epinasty, leaf bronzing discoloration starting from apex and lateral margins and progressing to mid-rib and base of the leaf) (Fig. S1). If any of such symptoms were observed—whether a few branches or most of the crown—the tree was considered positive for oak wilt. Note that the number of trees covered by each flight slightly varied due to different flight paths (Table S2). Current season crown wilt in 2017 suggested that crown wilt was present during mid to late August 2016 when airborne spectral data were collected. We georeferenced the canopy center of each tagged tree using a high-precision Trimble Pro6H GPS (Trimble, Sunnyvale, CA, USA) during the leaf-off stage the following winter 2017-2018. Finally, we georeferenced an additional 83 oaks (48 *Q. ellipsoidalis* and 35 *Q. macrocarpa*) that were not included in the training or testing steps of our models (see below). Instead, we used these oaks to further validate our models.

### 2.3 Canopy spectra extraction

We built a 1m-radius circular buffer around each canopy center using ArcGIS (version 10.6.1, ESRI, 2011) to sample fully sunlit canopy pixels per individual tree (Table S2), which were then linked to the respective species and oak wilt status (i.e., healthy, diseased). The number of pixels per tree ranged from four to nine—depending on the position of the buffer center in respect to that of the pixels within the buffer—for both AISA Eagle and AVIRIS-NG datasets with the AISA Eagle dataset yielding, on average, one to two more pixels per buffer due to its smaller pixel size. We applied the segmentation method designed to discard shaded canopy pixels and keep sunlit pixels only. We visually assessed reflectance in canopy pixels of a subset of trees for every species within our study and determined reflectance values near 552 nm, 671 nm, and 800 nm representative of sunlit canopy pixels (Malenovský et al., 2006). Then, we excluded pixels with values below such thresholds (0.01, 0.01, and 0.15 for 552 nm, 671 nm, and 800 nm, respectively) from our datasets. Spectral data processing employed the package *spectrolab* (Meireles et al. 2018) in R (version 3.6.0, R Development Core Team, 2020). First, we resampled the extracted spectral data to 410-980 nm for AISA Eagle and 410-2400 nm for AVIRIS-NG (both at 5 nm resolution to match wavelengths across sensors within the VNIR range) to remove noisy wavelengths at the range ends of the sensors and reduce the number of bands in the analyses. For AVIRIS-NG data only, we removed atmospheric water absorption bands between 1335-1430 nm and 1770-1965 nm and corrected artifacts at the sensor overlap region around 950 nm. Finally, for both datasets we unit vector-normalized reflectance values to reduce illumination differences among spectra (i.e., standardize differences in amplitude) (Feilhauer et al., 2010) while preserving differences in the shape of spectra that are important for species classification (Meireles et al., 2020b). After processing spectra, we calculated 21 spectral indices commonly used in the literature related to plant photosynthetic activity (e.g., RDVI, SIPI, SIF), water status (e.g., WBI, NDWI), and photoprotective stress (e.g., PRI, CRI700, NPQI) (see Table S1 for full index list). In cases where an index required a wavelength that was not a multiple of 5 and therefore missing in our spectra, we approximated the reflectance value of that wavelength based on the reflectance of the neighboring wavelengths either by using the nearest wavelength if the difference was ≤ 1 nm or otherwise by interpolation between the two nearest wavelengths. We assessed whether vector normalization affected the capacity of spectral indexes to detect oak wilt infected trees and found no major differences in spectral index performance (Appendix S1).

### 2.4 Statistical analyses

All statistical analyses were performed in R (version 3.6, R Development Core Team, 2020). To assess the capacity of canopy spectral reflectance to distinguish healthy trees from those infected with oak wilt, we performed partial least square discriminant analyses (PLS-DAs) (Barker and Rayens, 2003) using AISA Eagle (410-980 nm), AVIRIS-NG VNIR (410-980 nm), AVIRIS-NG SWIR (985-2400 nm) and AVIRIS-NG VSWIR (410-2400 nm). Performing PLS-DAs for each spectral range allowed us to assess the importance of each range of wavelengths for accurate detection. We treated each pixel as an observation because oak wilt disease does not manifest uniformly across the canopy of a tree, especially during early stages of infection. At early stages, the fungus may have infected only a fraction of the vessels within the tree trunk. Thus, curtailing the water supply to a few branches that become symptomatic while others remain asymptomatic. Treating pixels -rather than the whole tree- as observations is critical to prevent false negatives that result from early infected trees displaying a small number of symptomatic pixels. Thus, averaging pixels across a canopy composed of mostly healthy pixels may hide the signal from the infected pixels and lead to lower true positive classification rates as evidenced by the reduced performance of whole-tree level PLS-DAs aimed at distinguishing diseased from healthy trees (Fig. S2). In all PLS-DAs, we used ANOVA to compare models with different numbers of components and to identify the minimal number of components that maximized Kappa, a model performance statistic that quantifies model performance compared to random classification (Cohen, 1960). PLS-DAs were then run with the optimal number of components and the “Bayes” option to account for differences in prior probability distributions among classes (Brereton and Lloyd, 2014). The optimal number of components varied by model and are reported in the results section.

We tested the extent to which distinguishing red oaks from other species before oak wilt status classification improved the predictive performance of our models by evaluating two approaches for oak wilt detection: a modelling pipeline that did not consider species identities (“direct” approach) and one that differentiated red oaks from other species first (“phylogenetic” approach) (Fig. 1). Both approaches were applied to each sensor type and spectral range. In the direct approach, we ignored species identities and split the data within each class (“diseased” and “other”) into 75:25 randomly sampled subsets for model training and testing, respectively (Brereton and Lloyd, 2014; Fallon et al., 2020). We used the *caret* and *pls* packages in R (Kuhn 2008, Mevik et al. 2018) to assess model performance (accuracy, sensitivity, specificity, kappa) and obtain model-predicted values for each class (Congalton, 2001; Fassnacht et al., 2006). The random sampling, model training, model testing, performance assessment loop was iterated 10,000 times to generate 10,000 different training and test subsets, classification models, and corresponding performance estimates. We assessed overall performance of the direct approach by calculating the average and standard deviation of the performance outputs across all iterations.

**Figure 1:**
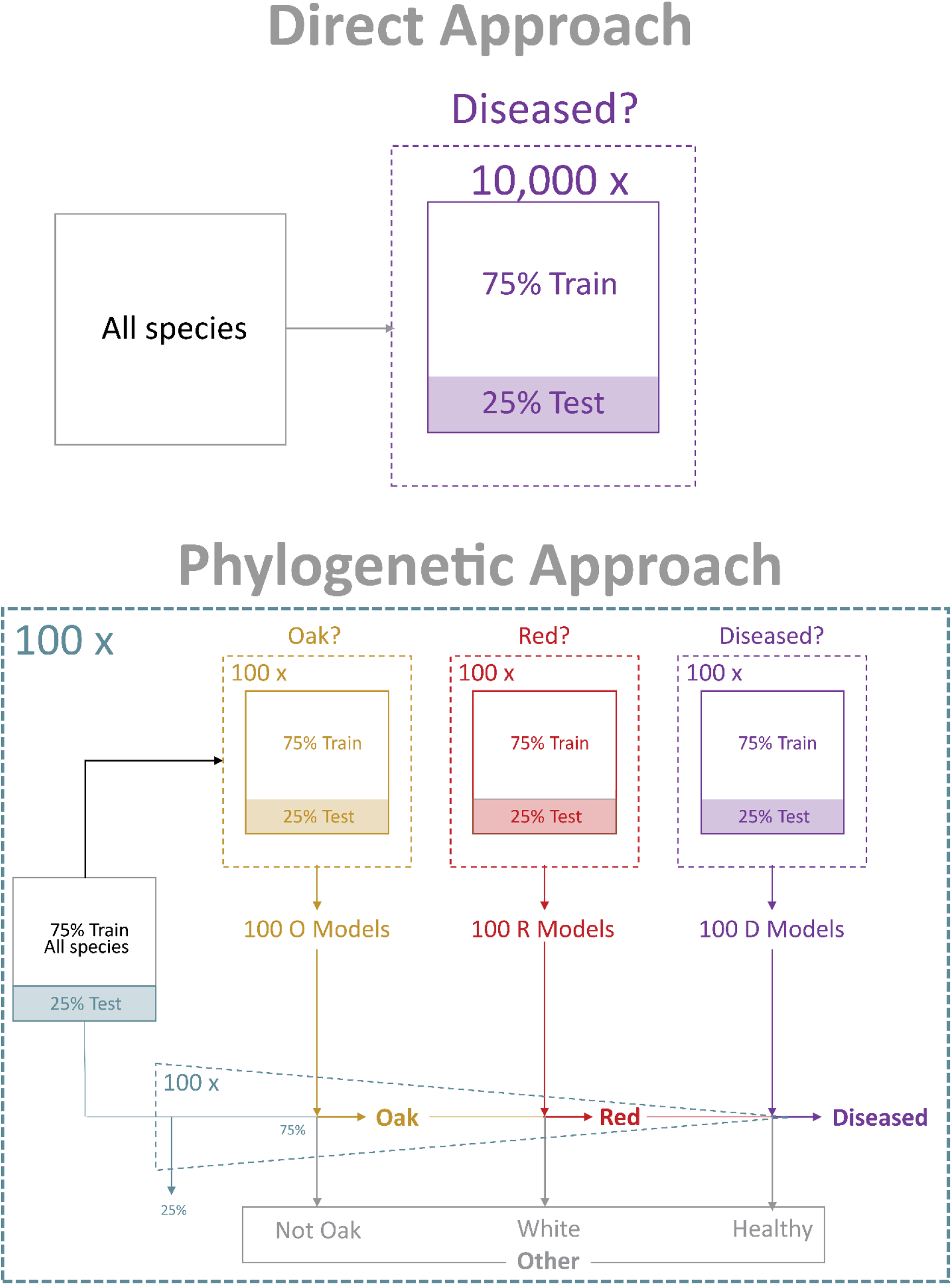
Workflow of the direct and phylogenetic modeling approaches used to classify diseased red oaks. In the phylogenetic approach, data were randomly split into 75% and 25% for model training and testing, respectively. The training set was used iteratively to train three sets of 100 models for distinguishing oaks from other species, red oaks from white oaks, and diseased red oaks from healthy red oaks. The trained models were coupled to filter out any observations that do not belong to the red oak group before running the disease detection step. This filtering process was tested using the initial 25% withheld test data. The whole process was iterated 100 times using different subsets of data to generate uncertainty around the performance estimates of the model. All classification results presented in the text utilize the 25% withheld data sets. See Table S3 for sample sizes within each step.

In the phylogenetic approach, we chained three distinctive PLS-DA types to solve the oak wilt classification problem sequentially through the steps illustrated in Fig. 1. First, we split our data into 75:25 randomly sampled subsets and left the 25% aside to test the overall performance of the phylogenetic approach at the end of the process (see below). Second, we used the 75% to train three types of PLS-DAs specifically aimed to distinguish 1) oaks from other species, 2) red oaks from white oaks, and 3) diseased red oaks from healthy red oaks. Accordingly, each model type had a different data structure: data from all species for PLS-DAs that distinguished oaks from other species, data belonging to the red and white oak group only for PLS-DAs that distinguished red from white oaks, and data including only putative red for PLS-DAs that distinguished diseased from healthy red oaks. All three PLS-DA types were performed following the same iterative approach described above by randomly sampling a subset of the 75% of the data for training, testing against the unused data of the subset (a 25% of the 75%), and assessing predictive performance of each PLS-DA. The purpose of these iterations was not to average model coefficients but rather to test how well PLS-DA types perform on average by generating confidence intervals for model performance estimates. We assessed performance of each PLS-DA type by calculating the average and standard deviation of the performance estimates across all iterations. We ran a total of 100 iterations for each PLS-DA type, thus obtaining 100 separate models of each type capable of distinguishing either oaks from other species, red oaks from white oaks, or diseased red oaks from healthy red oaks.

Finally, as an independent validation, we sequentially applied the 100 models of each PLS-DA type to the 25% of data originally set aside through another 100 iterations. During each iteration, the 25% subset containing all species was first split into 75:25 randomly sampled subsets (stratified by class, i.e., taxonomic grouping or health status) and only the 75% of the data were used with the aim of generating variation among iterations. In the first step of the phylogenetic pipeline, the selected data— which included all species —were classified as either oak or “other species” using the oak discrimination model. Then, the data classified as oak were classified as either “red” or “white oak” using the red oak discrimination model. Lastly, the data classified as red oak were classified as either “diseased” or “healthy red oak” using the disease discrimination model. Data classified as “other species”, “white oak”, or “healthy red oak” were later reclassified as “other” and their predicted classes were compared to their true identities to evaluate predictive performance. The full phylogenetic approach was iterated 100 times to ensure that the initial 75% split reflected all the existing variability within the dataset. Hence, we report performance across a total of 10,000 (100×100) models of each type (Fig. 1, see Table S3 for sample sizes and performance). We assessed overall performance of the phylogenetic approach by calculating the average and standard deviation of the multistep classification performance outputs across all iterations. Additionally, we applied the oak, and red oak PLS-DA models to the additional 83 oaks (48 *Q. ellipsoidalis* and 35 *Q. macrocarpa*) that were not included in any of the training or testing steps of our models as a second validation step.

Finally, we performed 100 direct PLS-DAs to classify the 12 dominant species present in our study area to identify those potentially causing misclassification of red oaks.

To determine which combination of wavelengths was most useful for detection of oak wilt, we extracted wavelength importance factors from PLS-DAs corresponding to AISA Eagle and AVIRIS-NG VSWIR and for both direct and phylogenetic approaches using the varImp() function in *caret* (Kuhn, 2008). We focused on these four PLS-DAs because they included the full range of wavelengths covered by each sensor with and without considering species identity. For simplicity, we limited our selection to the top 20 wavelengths with the highest average importance across all iterations within each model. We also extracted wavelength importance factors from AVIRIS-NG VNIR models to compare important wavelengths with the AISA Eagle models and identify shared wavelengths between both sensors. Lastly, we cross-applied AISA Eagle and AVIRIS-NG VNIR models to their respective testing datasets to assess the extent to which models developed with one sensor can be applied to data obtained from other sensors with similar range of wavelengths (i.e., sensor specificity).

To assess whether reflectance indices associated with physiology could distinguish healthy red oaks from those infected with oak wilt, we used ANOVA to perform pairwise comparisons between healthy and diseased red oaks across all spectral reflectance indices and for both AISA Eagle and AVIRIS-NG. Finally, we compared the effect sizes of these pairwise comparisons using Cohen’s *d* statistic (Cohen, 1988) to assess differences in the detectability of oak wilt between late July and late August.

### 2.5 PLS-DA mapping

As proof of concept, we mapped PLS-DA model outputs for predicted oak pixels, red oak pixels, and diseased red oak pixels across the landscape using an AISA Eagle flight line. We used AISA Eagle rather than AVIRIS-NG because the lower number of wavelengths significantly reduced the computational resources and time needed to generate the maps. First, we resampled and vector-normalized every pixel with the flight line using the same functions from the *spectrolab* package to match the spectral data used to train and test oak, red oak, and diseased red oak AISA Eagle PLS-DA models. For each model type (oak, red oak, and diseased red oak), we applied the coefficients of 100 randomly chosen model iterations of the 10,000 produced to each pixel of the resampled and normalized flight line using the function “predict” from package *car* (Fox and Weisberg, 2019). The result was 100 values per pixel— and thus 100 maps-describing the probability of being an oak, red oak, and a diseased red oak. We averaged the 100 values to obtain a single map with probability values representative of the average prediction. Then, we used a probability threshold of 0.5 to determine pixels with high certainty of being oaks, red oaks, and a 0.8 threshold for diseased red oaks to counter false positives resulting from the AISA model and generate masks to select pixels of the targeted classes. Lastly, we applied the masks to the original flight line both seen through true color and a combination of CMS, CI, and VOG2 indices in the red, green, and blue channels, respectively, to obtain landscape maps of oaks, red oaks, and diseased red oaks.

## 3. Results

All classification accuracy results are reported for the sets of 25% of samples withheld from the PLS-DA modeling steps, with the standard deviation calculated across the 10,000 iterations performed. All classification results are reported in Table S3.

### 3.1 Tree species identities can be predicted from canopy spectral reflectance

The tree species classification PLS-DA demonstrated that it is possible to accurately identify most of our 12 study species from spectral reflectance (AISA Eagle: 82.0% (±1.2%) correctly identified, AVIRIS-NG VSWIR: 90.0% (±1.3%), Appendix S2). Models correctly classified and differentiated white oaks (*Q. macrocarpa*) (AISA Eagle: 85.4% (±3.1%), AVIRIS-NG VSWIR: 94.7% (±1.3%)) and red oaks (*Q. ellipsoidalis* and *Q. rubra*) (AISA Eagle: 91.0% (±3.7%), AVIRIS-NG VSWIR: 99.4% (±1.4%)). However, models classifying the oak genus as a whole had higher accuracies (90.1% (±4.9%) and 98.6% (±2.9%) for AISA Eagle and AVIRIS-NG VSWIR, respectively, Appendix S3) than individual species models (Appendix S4), similar to results from leaf level spectra (Cavender-Bares et al., 2016). Accordingly, the validation steps also had higher performance when considering the oak genus as a whole (85.7% (±3.8%) and 94.7% (±2.5%) for AISA Eagle and AVIRIS-NG VSWIR, respectively, Appendix S3) instead of white and red oaks separately (59.9% (±3.7%) and 85.1% (±4.7%) for AISA Eagle and AVIRIS-NG VSWIR, respectively, Appendix S4).

### 3.2 Spectral reflectance models detected diseased red oaks

Spectral reflectance models did not accurately distinguish diseased red oaks from other trees unless red oaks were first distinguished from other species (Table S3). In the direct approach, overall model accuracy was significantly better than expected by chance (AISA Eagle: 58.9% (±5.0%), components (k) = 18; AVIRIS-NG VSWIR: 68.8% (±8.5%), k = 29, Fig. 2), but only healthy trees (true negatives) were correctly classified with high accuracy (AISA Eagle: 98.7% (±0.77%), AVIRIS-NG VSWIR: 97.5% (±1.4%), indicating high model specificity). Diseased red oaks were correctly classified (true positives) only 19.1% (±9.13%) and 40.0% (±15.5%) of the AISA Eagle and AVIRIS-NG VSWIR cases, respectively, indicating low model sensitivity (Fig. 2). As a result, isolating oaks and then red oaks through a stepwise phylogenetic PLS-DA model prior to disease detection increased correct classifications and improved the overall performance of both AISA Eagle and AVIRIS-NG VSWIR models (AISA Eagle: 75.4% (±6.4%), k = oaks: 16, red oaks: 10, diseased red oaks: 21; AVIRIS-NG VSWIR: 84.2% (±7.7%), k = oaks: 15, red oaks: 10, diseased red oaks: 16; Appendix S3-5, Table S3). The increase in performance was mostly due to a major increase in correct classification (true positives) of diseased red oaks (AISA Eagle: 54.6% (±11.5%), AVIRIS-NG VSWIR: 71.0% (±14.0%)) (Fig. 2, Appendix S5) resulting in increased model sensitivity compared to the direct PLS-DA approach.

**Figure 2:**
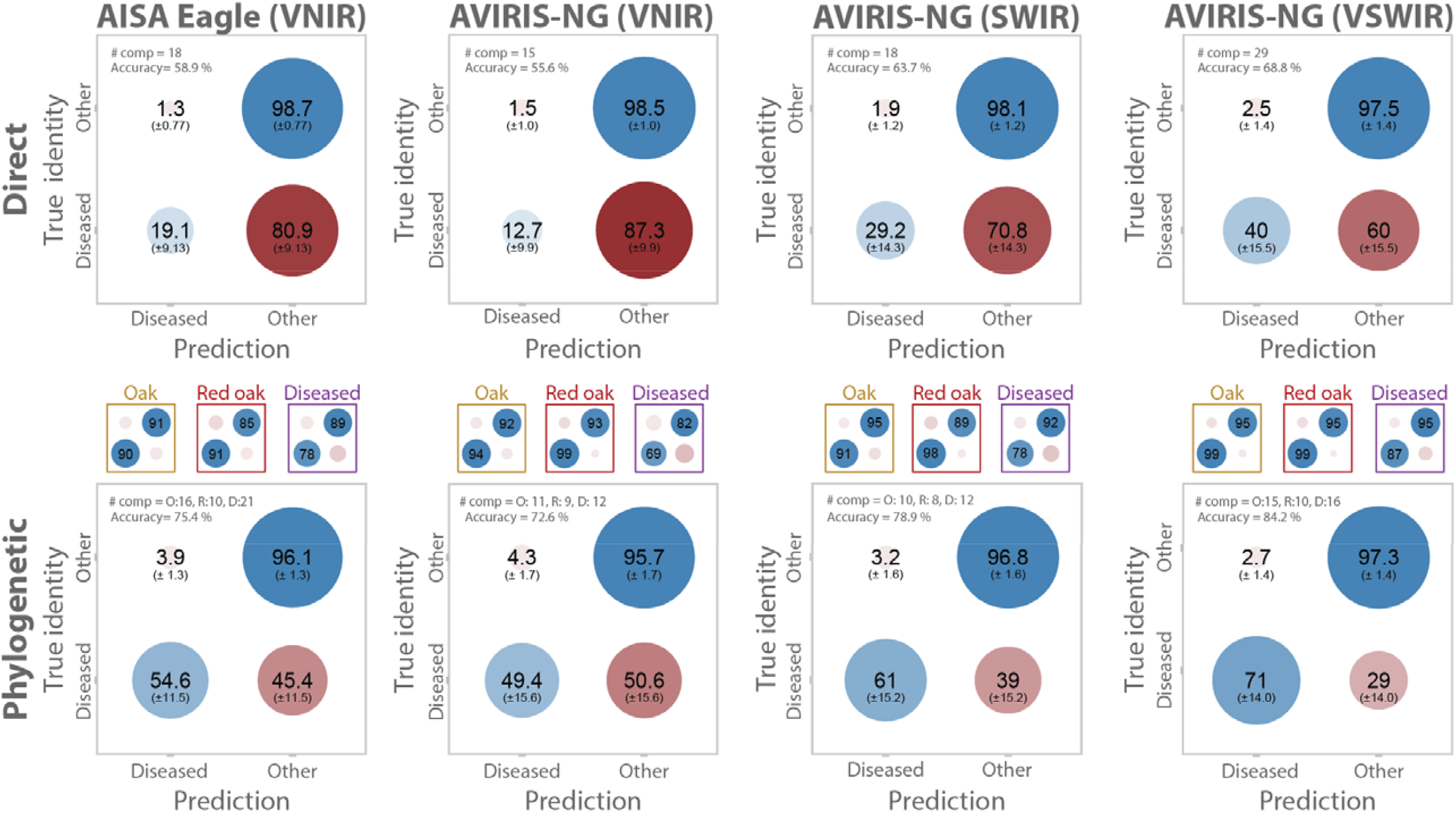
A stepwise phylogenetic classification approach enhanced detection of oak wilt in red oaks. Models that included both VNIR and SWIR wavelengths (AVIRIS-NG VSWIR) showed better prediction capacity than models including VNIR only. The short-wave infrared (SWIR) range is responsible for the increased predictive performance of AVIRIS-NG VSWIR relative to AVIRIS-NG VNIR PLS-DA models in the direct and phylogenetic modelling approaches. Blue and red circles represent correct and incorrect classifications, respectively. The size and color intensity of the circle represent the average percentage of classifications into each group based on the 25% of data withheld from 10,000 model-fitting iterations, one standard deviation is shown in parentheses. Grey boxes describe the overall predictive performance for a given approach and dataset. Red and blue circles in colored inset boxes above each phylogenetic model describe the performance of the steps within the phylogenetic model at discriminating oaks (gold), red oaks (red), and diseased red oaks (purple), respectively. The number of components used for each model or model step (O = oaks, R = red oaks, D = diseased) is given at the top left corner of the plot. See Appendices S3, S4, and S5 and Table S3 for detailed performance of the phylogenetic steps).

All steps within the phylogenetic PLS-DA model showed high performance (Table S3). The oak detection step showed high accuracy (AISA Eagle: 91% (±5.55%), k = 16; AVIRIS-NG VSWIR: 97% (±6.5%), k = 15) and correctly classified oaks in 90.7% (±6.2%) and 95.1% (±3.6%) of the AISA Eagle and AVIRIS-NG VSWIR cases, respectively (Appendix S3). Similarly, the red oak detection step showed high accuracy (AISA Eagle: 88% (±3.4%), k = 10; AVIRIS-NG VSWIR: 97% (±1.35%), k = 10) and correctly classified red oaks in 91.0% (±3.7%) and 99.4% (±1.4%) of the AISA Eagle and AVIRIS-NG VSWIR cases, respectively (Appendix S4). Finally, the diseased red oak detection step also showed high accuracy (AISA Eagle: 84% (±3.2%), k = 21; AVIRIS-NG VSWIR: 91% (±3.45%), k = 16) and correctly classified diseased red oaks in 78.4% (±3.1%) and 86.5% (±3.4%) of the AISA Eagle and AVIRIS-NG VSWIR cases, respectively (Appendix S5). The complexity of the model in this last step was higher in AISA Eagle (k = 21) than in AVIRIS-NG VSWIR models (k = 16). We note, however, that models were sensor-specific. AISA Eagle models failed to correctly distinguish classes when challenged with AVIRIS-NG VNIR data (Table S4). Similarly, AVIRIS-NG VNIR models failed to correctly distinguish classes when challenged with AISA Eagle data (Table S4).

### 3.3 VNIR and SWIR ranges are both important in detecting oak wilt

AVIRIS-NG SWIR models showed slightly higher classification accuracy (true positive rate) of diseased trees than AISA Eagle and AVIRIS-NG VNIR models in both direct (AVIRIS-NG SWIR: 63.7% (±7.75%), AISA Eagle: 58.9% (±5.0%), AVIRIS-NG VNIR: 55.6% (±5.45%)) and phylogenetic approaches (AVIRIS-NG SWIR: 78.9% (±8.4%), AISA Eagle: 75.4% (±6.4%), AVIRIS-NG VNIR: 72.6% (±8.65%)) (Fig. 2). When both AVIRIS-NG VNIR and SWIR were used together, models outperformed those using either VNIR or SWIR only. This was the case under both direct (AVIRIS-NG VSWIR: 68.8% (±8.45)) and phylogenetic approaches (AVIRIS-NG VSWIR: 84.2% (±7.7)).

When differentiating oaks from other species using AISA Eagle models, important wavelengths were clustered within the 500-560 nm, 600-620 nm, 660-690 nm, and the 970-980 nm regions of the VNIR range (Fig. 3). From these, wavelengths located around 550 nm and 675 nm where also important in AVIRIS-NG VNIR models. However, in AVIRIS-NG models that included both VNIR and SWIR, the importance of these regions was outweighed by regions 1200-1300 nm, 1440 nm, 1600-1750 nm, and 2230-2400 nm within the SWIR range. When differentiating red oaks from white oaks using AISA Eagle models, we found important wavelengths clustered within the 400-425 nm, 700-770 nm, and 925-980 nm regions. From these, wavelengths located at 420 nm and around 725 nm and 940 nm where also important in AVIRIS-NG VNIR models. In AVIRIS-NG VSWIR models, the importance of these VNIR regions remained high, but several regions within the SWIR range showed similar degree of importance. Within the SWIR, the important wavelengths were clustered at 1200 nm, 1440 nm, around 1490-1550 nm, and around 1700 nm. When differentiating healthy red oaks from diseased red oaks using AISA Eagle models, important wavelengths appeared at 725-750 nm and across the 800-980 nm region of the VNIR range. Wavelengths located around 725 nm, at 810 nm, 860, around 950 nm, and at 980 nm where also important in AVIRIS-NG VNIR models. AVIRIS-NG VSWIR models also identified as important wavelengths around 810 nm and 970 nm, but also identified important wavelengths within the SWIR range such as wavelengths around 1160 nm, 1260 nm, 1600 to 1750 nm, 1975 to 2050 nm and 2350 to 2400 nm. Most importantly, the twenty most important wavelengths for detection of oaks, red oaks, and diseased red oaks rarely overlapped in the AVIRIS-NG VSWIR models (Fig. 3). This was not the case for AISA Eagle models where most of the important wavelengths for red oak detection were the same than for diseased red oak detection.

**Figure 3:**
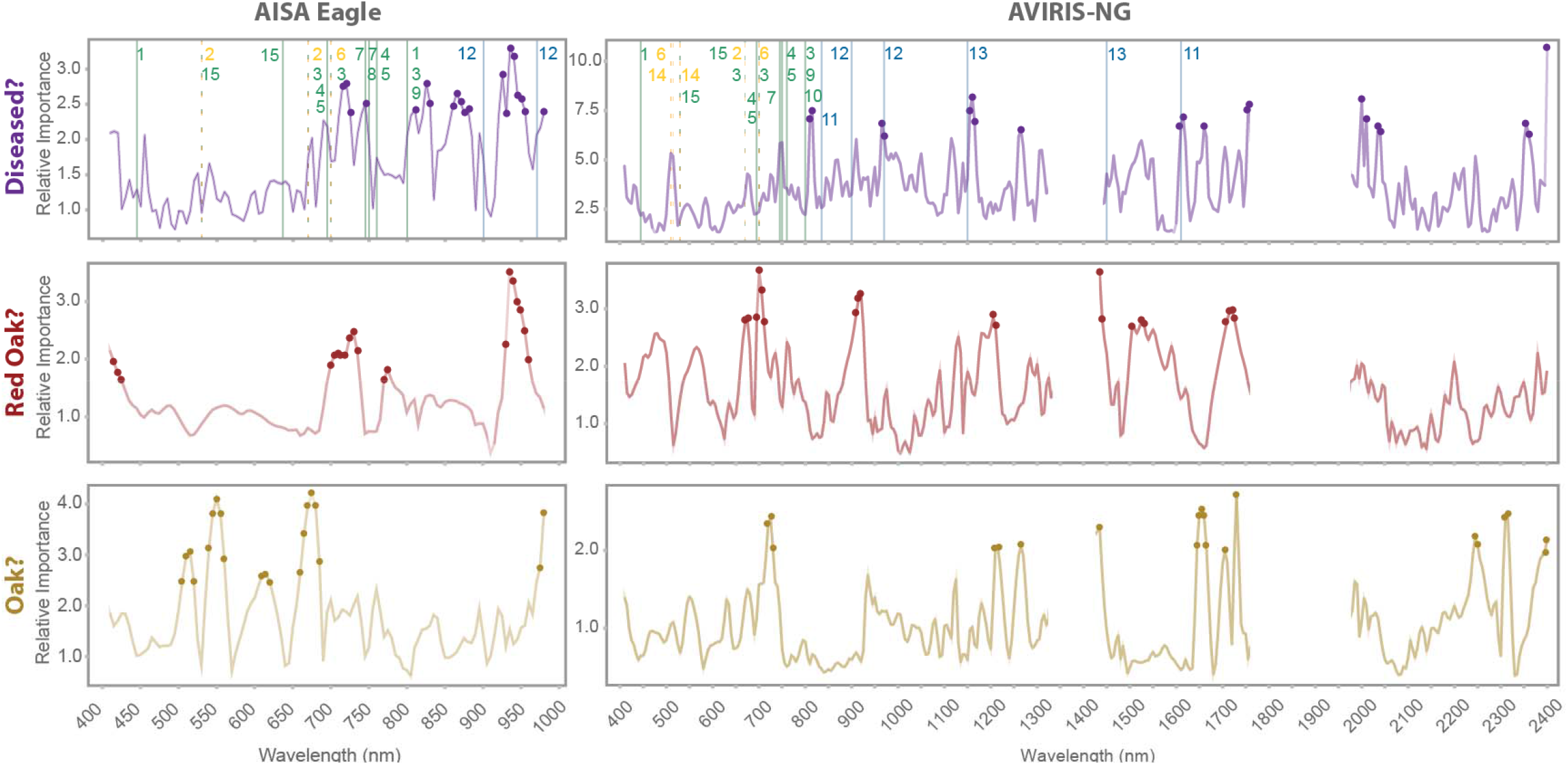
The twenty most important wavelengths—based on variable importance in projection (VIP)—differed among steps discriminating oaks (gold) from other species, red oaks (red) from white oaks, and diseased red oaks (purple) from healthy red oaks, and among models using either VNIR range (AISA Eagle) or both VNIR and SWIR ranges (AVIRIS-NG VSWIR). Vertical lines with numbers indicate wavelengths used in spectral indices associated with photosynthetic capacity (green), photoprotective pigment content (yellow), and water status (blue) that showed significant differences between healthy and oak wilt-infected trees. Numbers indicate spectral indices SIPI (1), PRIM4 (2), TCARI/OSAVI (3), CMS (4), SR_SIF_ (5), CRI700 (6), VOG2 (7), CI (8), RDVI (9), SR (10), NDWI (11), WBI (12), WBI-SWIR (13), PRIm1 (14), CCI (15).

### 3.4 Declines in photosynthetic capacity and water status signal oak wilt

Overall, spectral indices calculated from the AISA Eagle dataset collected during July showed slightly more pronounced differences between healthy and diseased red oaks than those calculated from the AVIRIS-NG dataset collected in August (Fig. 4). However, confidence intervals overlapped between both sensors for all photosynthetic and water status indices. Spectral indices associated with canopy photosynthetic capacity showed significant differences between healthy and diseased red oaks for both sensors and time periods. Within the AISA Eagle dataset, all indices associated with photosynthetic capacity were significantly different between healthy and diseased red oaks (Table S4). Within the AVIRIS-NG dataset, all indices associated with photosynthetic capacity except NPCI were significantly different between healthy and diseased red oaks. Most spectral indices associated with photoprotective stress only showed significant differences between healthy and diseased red oaks during the earliest flight (AISA Eagle). Those that were always significant—Photochemical Reflectance Index (PRIm4) and Carotenoid Reflectance Index (CRI700)—share wavelengths with indices of photosynthetic capacity, such as the SR and Transformed Chlorophyl Absorption in Reflectance Index/Optimized Soil-Adjusted Vegetation Index (TCARI/OSAVI) indices. Indices associated with canopy water status also showed significant differences between healthy and diseased red oaks. The effect sizes of the differences were comparable to those of indices associated with photosynthetic capacity.

**Figure 4:**
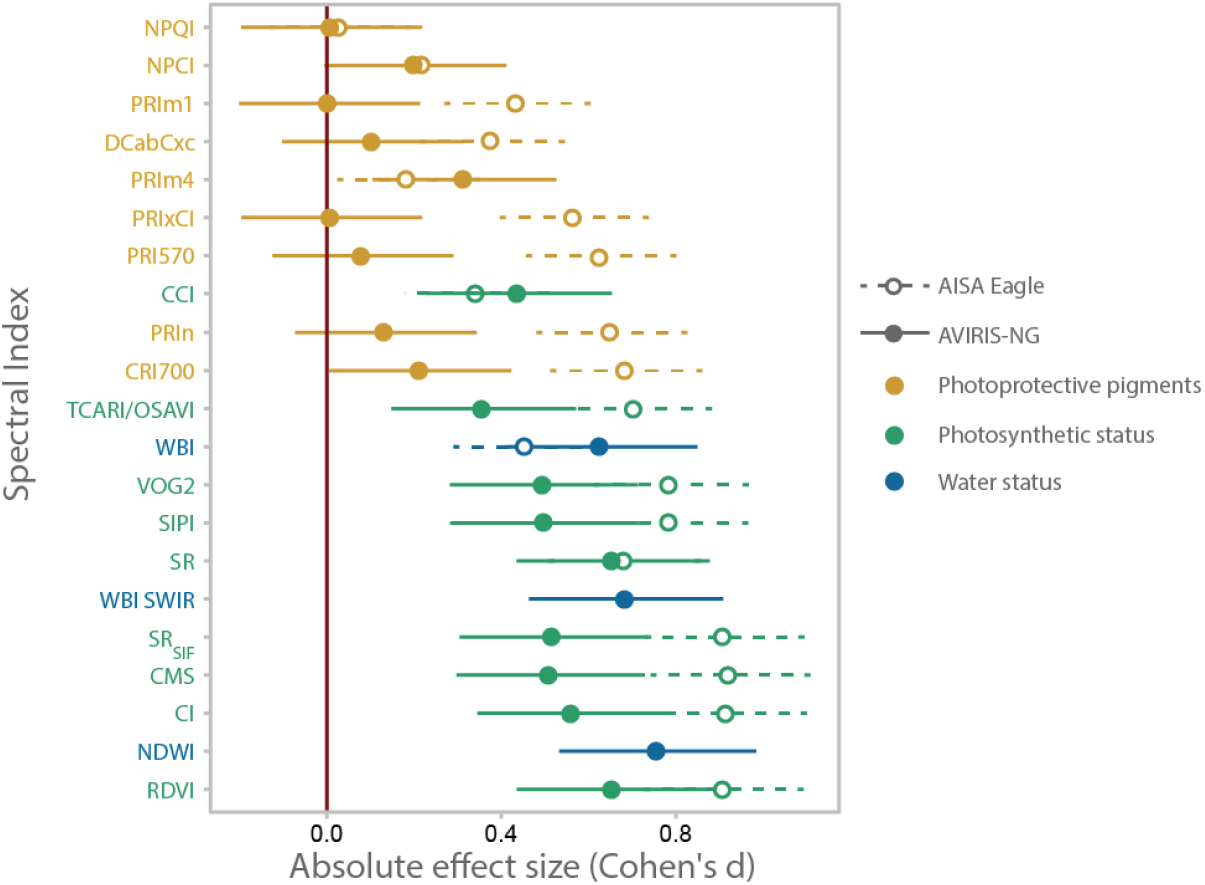
Spectral indices associated with photosynthetic (green) and water status (blue) differentiated diseased red oaks from healthy red oaks. Indices associated with photoprotective pigments (gold) only do so when collected in July or when they also include wavelengths associated with photosynthetic capacity. Each point represents the magnitude of the difference between healthy and diseased trees— shown by the absolute value of the Cohen’s d—for a given index and sensor type (AISA Eagle collected in July or AVIRIS-NG collected in August). Effect size can be understood as the amount of overlap between the distributions of two groups. For an effect size of 0, the mean of group 2 falls within the 50th percentile of group 1, and the distributions overlap completely, meaning there is no difference between them. For an effect size of 0.8, the mean of group 2 falls within the 79th percentile of group 1; thus, an average sample from group 2 would have a higher value than 79% of all samples from group 1 (Sullivan and Feinn, 2012). Lines represent 95% confidence intervals. Effect sizes are significantly different from zero when their confidence intervals do not overlap with the red zero line.

## 4. Discussion

The negative impacts of oak wilt and its rate of spread across North American ecosystems calls for detection tools that accurately identify trees affected by oak wilt at landscape scales (Haight et al., 2011; Hulcr and Dunn, 2011; Juzwik et al., 2011). We show that PLS-DA models developed from airborne spectroscopic imagery can detect oak wilt-infected red oaks. We demonstrate an approach to identify oak wilt-infected red oaks, which takes advantage of the physiological and phylogenetic information embedded in their reflectance spectra (Cavender-Bares et al., 2016; Meireles et al., 2020a). By first differentiating oaks from non-oaks, and then identifying red oaks—which are highly susceptible to rapid disease development—classification models based on spectral reflectance data can be used to distinguish oak-wilt affected and healthy red oaks with high accuracy. We also found that spectral indices associated with plant photosynthesis, water status, and photoprotective pigments are potentially sensitive to disease progression through physiological decline. Spectral indices also provide a mechanistic basis for understanding and tracking the physiological changes that allow classification models to detect oak wilt and are consistent with findings from other oak decline systems (Encinas-Valero et al. 2021).

### 4.1 Including short wave infrared reflectance improves model accuracy

Including SWIR wavelengths in spectral reflectance models increases oak wilt detection accuracy. We observed higher oak wilt detectability in direct AVIRIS-NG SWIR and VSWIR than direct AISA Eagle and AVIRIS-NG VNIR models. Direct PLS-DAs using AVIRIS-NG VNIR showed similar performance to that of AISA Eagle VNIR models (Fig. 2). We can therefore attribute the greater performance of direct AVIRIS-NG VSWIR models to the addition of SWIR wavelengths. Direct AISA Eagle models rely on many of the same wavelengths to distinguish red oaks from white oaks and to distinguish diseased and healthy red oaks (Fig. 3). As such, they often misclassify diseased red oaks as white oaks (Fig. 2, Appendix S4). However, even direct models show much higher accuracy when both VNIR and SWIR ranges are included (AVIRIS-NG VSWIR). The additional information-rich SWIR wavelengths allow models to use different wavelength regions to distinguish oaks from other species, red oaks from white oaks, and diseased red oaks from healthy red oaks (Fig. 2). Consequently, the critical wavelengths to identify oaks, red oaks, and diseased red oaks overlap less, which reduces the chances of confusion among classes (Fig. 3 and validation steps in Appendices S3 & 4). Most likely, including SWIR reflectance provides temporally stable spectral features containing phylogenetic information associated with plant structural traits (Cavender-Bares et al., 2020; Meireles et al., 2020a) that serve to reduce misclassification of diseased red oaks as white oaks and other species. Indeed, we find that the SWIR wavelengths were more often represented than VNIR wavelengths in oak models and as represented as VNIR wavelengths in red oak models (Fig. 3). In particular, 17 and 11 of the most important wavelengths for identifying oaks and red oaks, respectively, fell within the SWIR range. The SWIR was also important for distinguishing diseased from healthy red oaks. Among the most important SWIR wavelengths were those associated to plant water content and leaf chemistry such as protein, sugars, lignin, and cellulose content (Asner et al., 2018; Fourty et al., 1996). Based on our results, the SWIR range appears to contain disease-specific and phylogenetic information highly relevant to detecting symptoms of oak wilt and to identifying its hosts. Hence, similar to previous work combining VNIR reflectance with SIF or thermal data (Zarco-Tejada et al., 2018, 2016), when SWIR wavelengths are combined with VNIR in oak wilt detection models, detection rates are maximized.

### 4.2 A multi-step phylogenetic approach increases accuracy

Partitioning the classification process into simple binary steps within a phylogenetic framework reduces potential misclassification and increases model accuracy. We used a hierarchical classification approach (Allen and Walsh, 1996; Townsend et al., 2009; Wolter et al., 1995) aimed towards distinguishing the more susceptible red oaks from white oaks and other species that are less susceptible to oak wilt. During the first step, phylogenetic models distinguish between oaks and other species because the reflectance spectrum shows phylogenetic conservatism among the oaks (Cavender-Bares et al., 2016; Cavender-Bares, 2019), including those infected by oak wilt. The model is not required to distinguish between healthy and diseased conspecifics in this first step, thus simplifying the task. Reducing the number of potential classes becomes increasingly important as the individuals become more phylogenetically related—and hence more phenotypically similar—which makes correct classification more challenging (Meireles et al., 2020b). Removing non-oak species significantly reduces variation in phylogenetically conserved regions of the spectra, allowing the model to be trained on spectral differences that distinguish white and red oaks and subsequently on the spectral variation that distinguishes diseased and healthy red oaks. Because of these filtering steps, the disease detection algorithm is highly accurate (84%; Fig. 2, appendix S5) and significantly more accurate than a single-step, direct approach. While the phylogenetic approach gains complexity in terms of number of steps, each binary classification step is simple and requires few independent components. Although each step generates classification errors that propagate through the modeling pipeline, these errors are captured by the overall performance metrics, indicating that the increase in accuracy gained through the phylogenetic filtering outweighs the propagated errors. The phylogenetic approach increases accuracy by reducing the number of classes to compare while inclusion of the SWIR range increases accuracy by increasing the number of informative wavelengths.

While the PLS-DA models are sensor-specific (Table S4), the benefits of implementing a stepwise phylogenetic approach or using the widest range of wavelengths possible do seem common across hyperspectral products. Implementing a stepwise phylogenetic approach boosted model performance to a similar extent in models that had the same range of wavelengths (i.e., AISA Eagle and AVIRIS-NG VNIR models). Similarly, reducing the range of wavelengths in AVIRIS-NG models to the VNIR range reduced model performance to similar values observed in AISA Eagle models. The similar performance between datasets from different sensors and time periods could suggest that the benefits of adding SWIR wavelengths and a stepwise phylogenetic approach are insensitive to sensor type or time of the year. Furthermore, despite the sensor-specificity of the models, several wavelengths were commonly highlighted as important across sensor types in oak, red oak, and diseased red oak models (Figure S3, see results section). These wavelengths were located around the green band and close to the red edge in case of the oak model. In the case of the red oak and diseased red oak models, shared wavelengths were also close to the red edge, and representative of starch and water content (Curran, 1989).

Our results highlight that species classification is critical for increasing model accuracy for a simple reason: if the disease is host-specific, modeling can be more tractable by detecting potential hosts first. Future studies should test whether phylogenetic models with simple binary classification steps such as the one used here make disease detection models generic enough to be applicable across different sites and years.

### 4.3 Targeted spectral indices help understand physiological changes associated with oak wilt disease

Diseased red oaks were more easily differentiated from healthy red oaks by spectral reflectance indices associated with photosynthetic activity (Carter and Knapp, 2001; Vogelmann et al., 1993; Zarco-Tejada et al., 2002) and water status (Ceccato et al., 2001; Serrano et al., 2000; Ullah et al., 2014) than by photoprotective pigment content indices. Indices based on photoprotective pigment only differentiated diseased trees from asymptomatic trees in July or when they included wavelengths also associated with photosynthetic activity (Figs. 3 & 4). These results suggest that oak wilt infection in adult trees in natural ecosystems first triggers changes in photoprotective pigments (similar to holm oak decline caused by *Phytopthora spp* fungal infection and drought, Encinas-Valero et al. 2021) followed by declines in photosynthetic rate, stomatal conductance, and water content just as in greenhouse red oak seedlings infected with oak wilt (Fallon et al., 2020). Our results uncover an important temporal pattern in physiological decline: photoprotective pigment content ceases to be a good predictor of oak wilt by the end of summer. By July, indices of photoprotective pigment content showed somewhat similar sensitivity to oak wilt than indices of photosynthetic capacity or water status (Fig. 4). At early stages, both tylose production (in response to infection) and plugging of vessels by metabolites of the fungus are likely to have contributed to diminished water transport. In turn, reduced transpiration and stomatal closure induced by reduced water supply is expected to have caused photosynthetic decline (Fallon et al., 2020) and abnormally increased photoprotective pigment content to deal with light stress resulting from reduced photosynthesis. By August, healthy trees may also have increased photoprotective pigment content as summer drought becomes more prevalent and photosynthesis is partially impaired. Yet, diseased trees have likely experienced greater water deficit due to tylose formation and their photosynthetic capacity and water status is impaired to a greater extent than drought-stressed but uninfected trees. This would explain why photosynthetic capacity indices still distinguished diseased and healthy red oaks and why water status indices showed greater overall sensitivity to oak wilt than photosynthetic activity indices by August (Fig. 4). The fact that, photosynthetic and water status indices showed similar performance between sensors suggests that normalized spectral indices should be comparable and applicable across sensor types to some degree due to their underlying mechanistic basis. The results are consistent with experimental work indicating that photosynthesis is the first physiological process to decline as stomata shut down (Fallon et al., 2020) followed by water content as vessel occlusion develops and the fungus damages cell walls and membranes (e.g., through pathogen-produced toxins) leading to dehydration and tissue death (Oliva et al., 2014). While other studies have proposed the use of machine learning algorithms to identify tree species and the use of spectral indices to detect plant diseases (Abdulridha et al., 2019; Ghosh et al., 2014; Iordache et al., 2020; Pontius, 2014; Tang et al., 2021), we see potential in coupling both. Pairing phylogenetic approaches to map oak wilt presence with photosynthetic activity and water status indices can provide powerful tools to delineate oak wilt centers across areas of the landscape. The steps would entail first mapping red oaks (Fig. S4) - and diseased red oaks if possible- and then combining spectral indices associated with photosynthetic and water status, or indices that use oak-wilt sensitive wavelengths to detect potential oak wilt pockets (Fig. 5). Pockets of affected trees may show a center-outward radial gradient with dry dead trees at the center, dehydrated and photosynthetically impaired trees in the middle, and trees with slightly lower photosynthetic capacity than expected around the edge of the pocket (i.e., early disease development phase) (Figs. 5, S4, S5 & S5). The step of distinguishing oaks and red oaks from other species across the landscape is highly advantageous for forest managers on its own as it allows them to identify areas at high risk of infection in which to apply spectral indices for further diagnostic. Hence, phylogenetic spectral models paired with indices is a novel approach to predict the risk of disease spread based on the abundance of red oaks across the landscape and the stage of disease development based on physiological status, thus allowing managers to better assess risk of spread and adjust the magnitude of their interventions accordingly (Pontius and Hallett, 2014).

**Figure 5.**
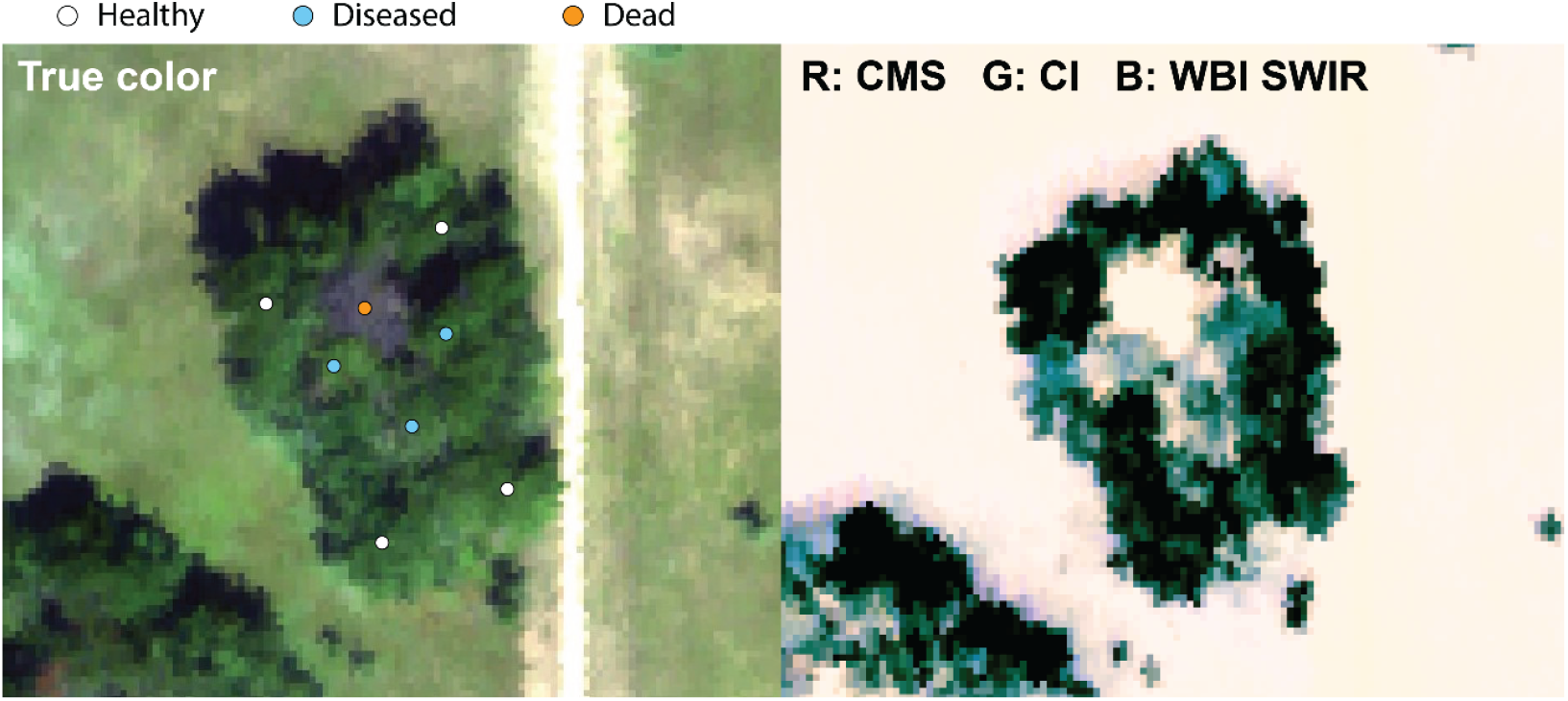
A typical oak wilt pocket observed through true color and a combination of spectral indices using the 2016 AVIRIS-NG data. A tree killed by oak wilt during 2015 (orange circle) can be observed at the center of the oak wilt pocket in true color (red as 640 nm, green as 550 nm, and blue as 470 nm). Three diseased trees (blue circles) stand next to it that cannot be detected with true color images. Both dead and diseased trees are surrounded by an outer ring of healthy trees. Diseased trees become apparent through spectral indices associated with photosynthetic function and water status -such as the Carter-Miller Stress index (CMS), Chlorophyll Index red edge (CI), and Water Band Index in the SWIR range (WBI SWIR)-placed on the red (R), green (G), and blue (B) channels.

## 5. Conclusions

Remote detection greatly enhances the ability of managers to prevent the enormous ecological and economical damage caused by invasive species (Juzwik, 2000; Poland et al., 2021). Airborne spectroscopic imagery enables landscape-level detection of diseases caused by invasive pathogens, like oak wilt, due to the phylogenetic and physiological information embedded in spectral reflectance. SWIR wavelengths increased model accuracy by enabling detection of disease-specific hosts, a critical step in identifying forested areas vulnerable to infection. Additionally, inference of the physiological basis of oak wilt symptom development using spectral indices associated with known spectral features points to the potential to delineate oak wilt centers using remote sensing products that monitor canopy photosynthetic capacity and water status such as VOG2, SIPI, SR_SIF_, CMS, CI, RDVI, WBI, and NDWI among others. Importantly, in our study landscape detection was made possible by coupling airborne spectroscopic imagery with traditional knowledge from taxonomic and disease experts and high precision ground GPS reference data. While landscape detection of oak wilt will facilitate the task of detecting infected trees, there is still much work to do. Future studies should assess whether PLS-DA models will be general enough to detect oak wilt across years and sites and whether the physiological basis of oak wilt symptom development will be sufficient to make accurate inferences about the presence of new oak wilt infections. Further investigation of the physiological changes that accompany disease progression may also provide the link to scale spectral detection to regional scales via spaceborne platforms. The work done here points to the benefit of research that might lead to an “optimal” remote sensing system (airborne or satellite) for detecting invasive diseases. We hope that our research motivates such work.

## Acknowledgements

This project was funded by the Minnesota Invasive Terrestrial Plants and Pests Center, the National Science Foundation (NSF) and the National Aeronautics and Space Administration (NASA) through the Dimensions of Biodiversity program (DEB-1342872 and DEB-1342778), the Cedar Creek Long Term Ecological Research program (1831944), and the NSF Biology Integration Institute ASCEND (DBI: 2021898). The University of Minnesota, including Cedar Creek ESR, lies on the ancestral, traditional, and contemporary Land of the Dakota people. We would like to thank Brett Fredericksen, Erin Murdock, Kali Hall, Lewis French and Travis Cobb for help with tarp measurement for empirical line corrections. Thanks to Dr. Joe Knight for access to the infrastructure of the Remote Sensing and Geospatial Analysis Lab at UMN, and to Dan Bahauddin for informatics help which facilitated flight planning at Cedar Creek. We thank Paul Castillo, USFS for his technical assistance in the training of field technicians, and Zhihui Wang for assistance processing AVIRIS-NG imagery. We also thank Shan Kothari, Lucy Shroeder, Anna Yang, Austin Yantes, Clarissa Fontes, Byju Govidan, Artur Stefanski, and Adriana Castillo Castillo for their comments in previous versions of this manuscript.

## Notes

### Competing Interest Statement

The authors have declared no competing interest.

### Summary of Updates

Edits requested by reviewers integrated into the manuscript.

