## Appendix for "Canopy spectral reflectance detects oak wilt at the landscape scale using phylogenetic discrimination"

**Supplementary Information**

**Table S1:** List of spectral indices used and the physiological characteristics that they reflect. Numbers preceded by an “R” within formulae indicate reflectance wavelengths. In cases where an index required a wavelength that was not a multiple of 5 and therefore missing in our spectra, we approximated the reflectance value of that wavelength based on the reflectance of the neighboring wavelengths as detailed in the main text. References for each index are provided at the end of the table.

| Spectral Index | Physiological characteristic | Formula | Reference |
| --- | --- | --- | --- |
| *Carotenoid Reflectance Index 700 (CRI700)* | Carotenoid content | (1/R510) - (1/R700) | Gitelson et al. (2003; 2006) |
| *Vogelmann index (VOG2)* | Chlorophyll and water content | (R734 - R747)/(R715 + R726) | Vogelmann et al. (1993) |
| *Normalized Pigment Chlorophyl Index (NPCI)* | Chlorophyll; general stress | (R680 - R430)/(R680 + R430) | Peñuelas et al., (1995) |
| *Structure Insensitive Pigment Index (SIPI)* | Carotenoid : chlorophyll ratios | (R800 - R445)/(R800 + R680) | Peñuelas, Baret, and Filella (1995) |
| *Chlorophyll Index Red Edge (CI)* | Chlorophyll and chlorophyll fluorescence | (R750/R710) | Haboudane et al. (2002) |
| *Transformed Chlorophyl Absorption in Reflectance Index/Optimized Soil-Adjusted Vegetation Index (TCARI/OSAVI)* | Chlorophyll content corrected by canopy density | 3 * [(R700 - R670) - 0.2 * (R700 - R550) * (R700/R670)]/((1 + 0.16) * (R800 - R670)/(R800 + R670 + 0.16)) | Haboudane et al. (2002) |
| *Simple Ratio using the O2A SIF band (SR_SIF_)* | Chlorophyll content | R690/R760 | Freedman et al. (2002) |
| *Simple Ratio (SR)* | Leaf structural integrity | R800/R670 | Jordan (1969) |
| *Renormalized Difference Veg. Index (RDVI)* | Leaf structural integrity | (R800 - R670)/sqrt(R800 + R670) | Roujean & Breon (1995) |
| *Carter Miller Stress (CMS)* | Photosynthetic activity | R694/R760 | Carter (1994) |
| *Carotenoid/Chlorophyl ratio index (PRIxCI)* | Carotenoid : chlorophyll ratios | (R570 - R530)/(R570 + R530) * ((R760/R700) - 1) | Garrity et al. (2011) |
| *Chlorophyl/Carotenoid Index (CCI)* | Carotenoid : chlorophyll ratios | (R530 – R645)/ (R530 + R645) | Gamon et al. (2016) |
| *Water Band Index (WBI)* | Canopy water content | R970/R900 | Peñuelas et al., (1997) |
| *Normalized Difference Water Index (NDWI)* | Canopy water content | (R835 - R1610)/(R835 + R1610) | Gao (1996) |
| *Water Band Index in SWIR range (WBI_SWIR)* | Canopy water content | R1150/R1450 | New from this paper |
| *Normalized Photochemical reflectance index (PRIn)* | Xanthophyll cycle activity | PRI570/[RDVI * (R700/R670)] | Zarco-Tejada et al. (2013) |
| *Photochemical reflectance index 570 (PRI570)* | Xanthophyll cycle activity | (R570 - R531)/(R570 + R531) | Gamon et al. (1992) |
| *Normalized Phaeophytinization Index (NPQI)* | Chlorophyll; general stress | (R415 - R435)/(R415 + R435) | Peñuelas et al., (1995) |
| *Photochemical reflectance index 512 (PRIm1)* | Xanthophyll cycle activity | (R512 - R531)/(R512 + R531) | Hernández-Clemente et al. (2011) |
| *Photochemical reflectance index 670 & 570 (PRIm4)* | Xanthophyll cycle activity | (R570 - R531 - R670)/(R570 + R531 + R670) | Hernández-Clemente et al. (2011) |
| *Reflectance band ratio index (DCabCxc)* | Xanthophyll cycle activity | R672/(R550 * 3R708) | Datt et al. (1998) |

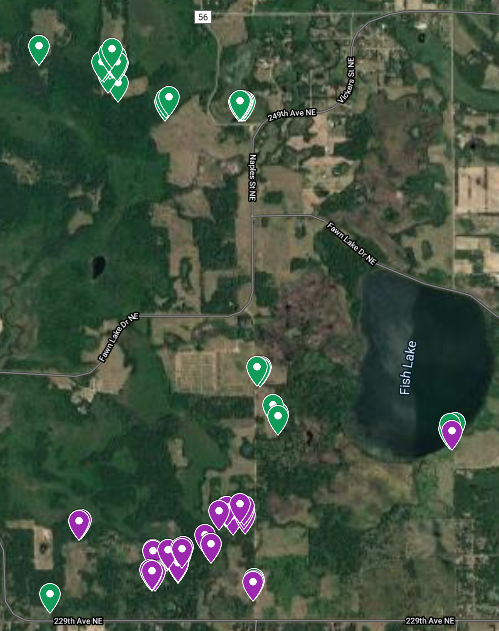

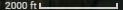

**Figure S1:** Aerial view of study area. Locations of the healthy (green) and diseased (purple) red oaks used in this study are shown.

**Table S2** List of species included in the study, the number of trees tagged for each species, and the number of sunlit and shaded pixels obtained from these trees using AISA Eagle and AVIRIS-NG spectroscopic imagery. Diameter at Breast Height (DBH) was calculated from circumference at breast height.

| **TRAINING & TESTING** | | | | | | | | | | | |
| --- | --- | --- | --- | --- | --- | --- | --- | --- | --- | --- | --- |
|  |  |  |  | **AISA Eagle** | | | | **AVIRIS-NG** | | | |
| **Species** | **ID** | **Height range (m)** | **DBH range (cm)** | **Trees** | **Sunlit and**  **shaded pixels** | **Sunlit**  **pixels** | **Shaded**  **pixels** | **Trees** | **Sunlit and**  **shaded pixels** | **Sunlit**  **pixels** | **Shaded**  **pixels** |
| *Acer rubrum* | ACRU | 11-22 | 9-42 | 8 | 52 | 50 | 2 | 6 | 24 | 24 | 0 |
| *Acer saccharinum* | ACSA | 13-25 | 61-190 | 15 | 93 | 74 | 19 | 15 | 63 | 62 | 1 |
| *Betula papifera* | BEPA | 9-24 | 11-34 | 34 | 212 | 203 | 9 | 31 | 131 | 131 | 0 |
| *Larix laricina* | LALA | 5-10 | 10-18 | 40 | 243 | 242 | 1 | 39 | 170 | 158 | 12 |
| *Pinus resinosa* | PIRE | 16-26 | 17-50 | 42 | 249 | 249 | 0 | 42 | 172 | 169 | 3 |
| *Pinus strobus* | PIST | 6-32 | 20-78 | 56 | 335 | 334 | 1 | 51 | 297 | 287 | 10 |
| *Pinus sylvestris* | PISY | 7-20 | 16-76 | 33 | 197 | 193 | 4 | 30 | 130 | 125 | 5 |
| *Populus grandidentata* | POGR | 15-25 | 19-51 | 40 | 239 | 239 | 0 | 39 | 235 | 235 | 0 |
| *Populus tremuloides* | POTR | 9-20 | 2-48 | 40 | 246 | 244 | 2 | 9 | 39 | 36 | 3 |
| *Quercus ellipsoidalis* | QUEL | 8-26 | 19-72 | 47 | 280 | 271 | 9 | 40 | 174 | 174 | 0 |
| *Quercus macrocarpa* | QUMA | 7-25 | 13-74 | 49 | 308 | 308 | 0 | 40 | 166 | 151 | 15 |
| *Quercus rubra* | QURU | 21-24 | 18-87 | 19 | 112 | 109 | 3 | 18 | 73 | 73 | 0 |
| *Quercus ellipsoidalis (infected)* | OWQU | 15-26 | 25-63 | 41 | 239 | 234 | 5 | 38 | 140 | 140 | 0 |
| **Total** | **TOTAL** |  |  | **464** | **2805** | **2750** | **55** | **398** | **1814** | **1765** | **49** |
| **VALIDATION** | | | | | | | | | | | |
| *Quercus ellipsoidalis* | QUEL | 8-26 | 19-72 | 48 | 292 | 288 | 4 | 49 | 185 | 185 | 0 |
| *Quercus macrocarpa* | QUMA | 7-25 | 13-74 | 35 | 233 | 219 | 14 | 47 | 181 | 180 | 1 |
| **Total** | **TOTAL** |  |  | **83** | **525** | **507** | **18** | **96** | **366** | **365** | **1** |

**Appendix S1** Vector normalization (standardization to unit length) was applied to spectra to standardize differences in amplitude while preserving differences in shape that are important for species classification. Panel A corresponds to mean spectra of AISA Eagle data while panel B corresponds to AVIRIS-NG data. Left columns represent spectra for all classes (for abbreviations see Table S2). Right columns emphasize the effects of vector normalization on healthy and diseased red oak mean spectra. Panel C demonstrates that spectral indices associated with photosynthetic function and water content were the most relevant for oak wilt detection regardless of whether vector normalization was applied. In panel C, each point represents the magnitude of the differences between healthy and diseased trees—shown by the absolute value of the Cohen’s d—for a given index and time of data collection (July or August). Lines represent 95% confidence intervals. Effect sizes are significantly different from zero when their confidence intervals do not overlap with the red zero line.

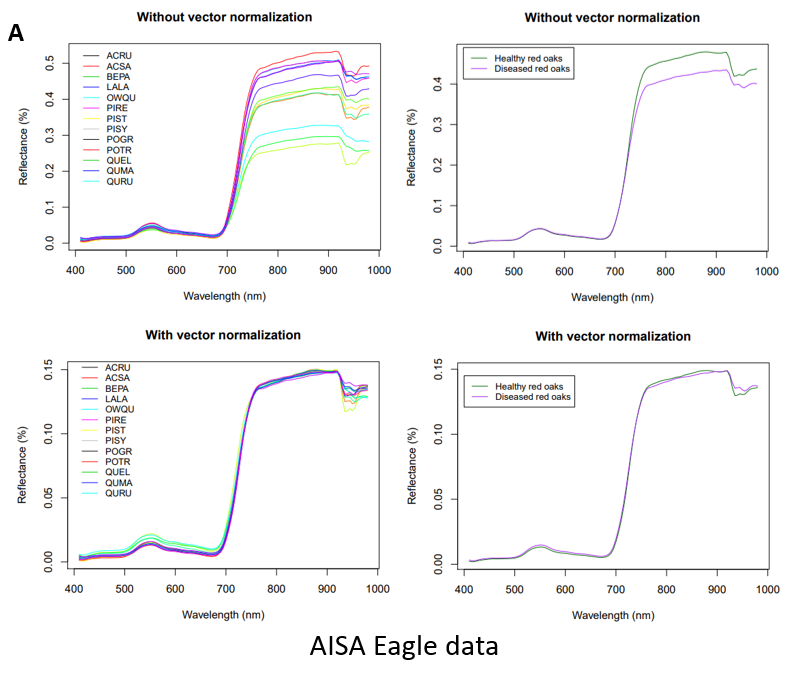

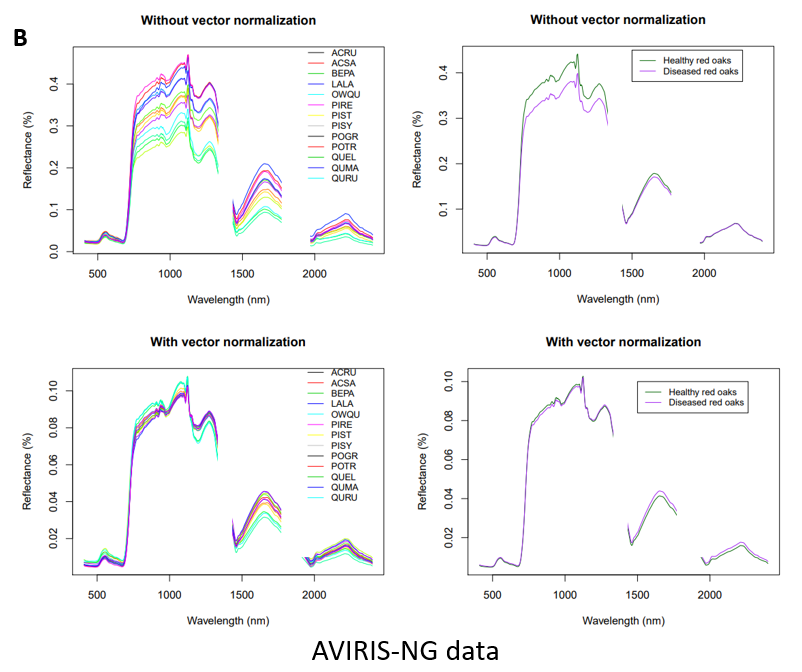

**C**

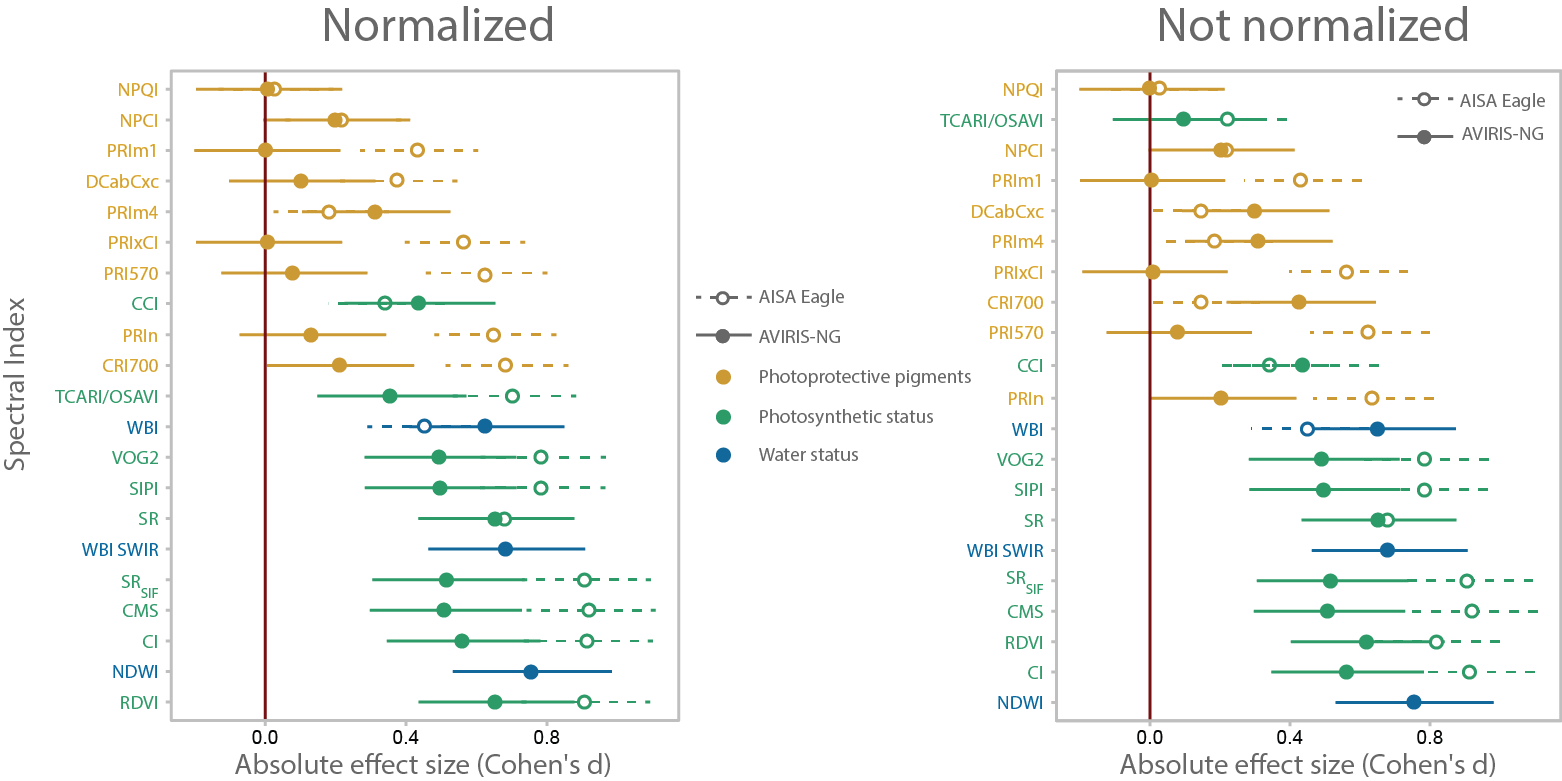

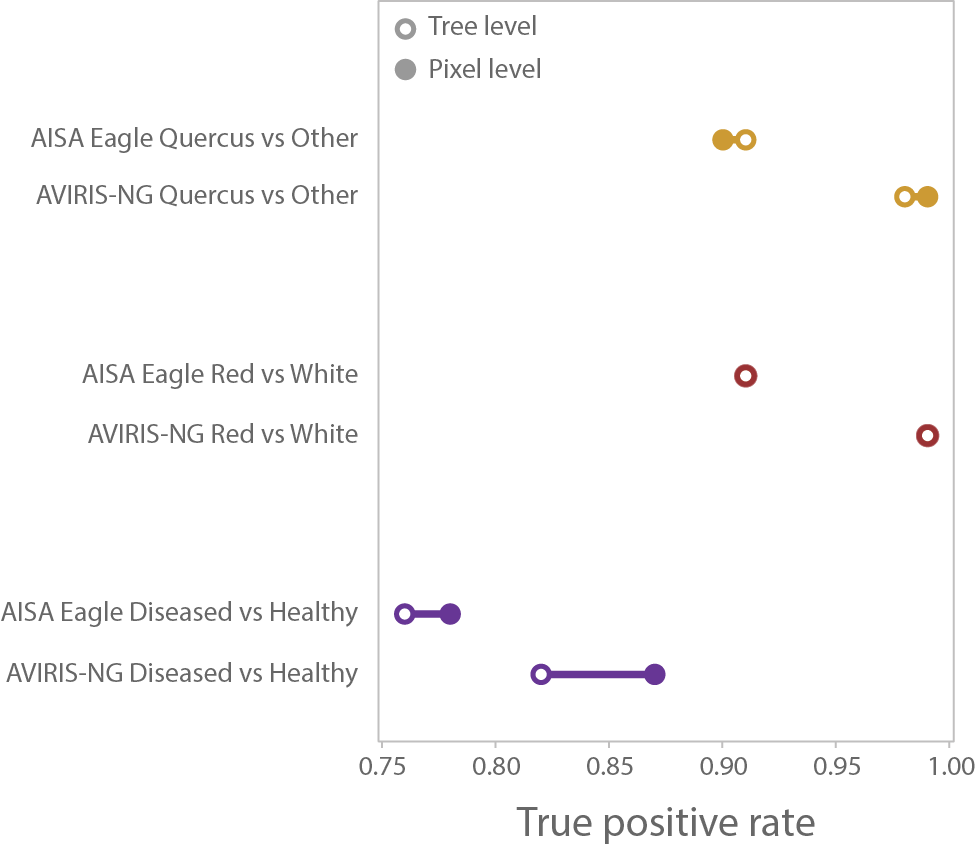

**Figure S2.** A tree-level approach reduces detectability of diseased red oaks because the oak wilt disease does not manifest uniformly across the canopy of a tree, especially during early stages of infection. At early stages, the fungus may have infected only a fraction of the vessels within the tree trunk. Thus, curtailing the water supply to a few branches that become symptomatic while others remain asymptomatic. Treating pixels -rather than the whole tree- as observations is critical to maximize early detection because early infected trees may display a small number of symptomatic pixels. Thus, averaging pixels across a canopy composed of mostly healthy pixels hides the signal from the infected pixels and leads to lower true positive classification rates.

**Appendix S2** Species identification based on AISA Eagle visible to near-infrared (VNIR) and AVIRIS-NG VNIR plus shortwave infrared (VSWIR) spectra. Blue and red circles represent correct and incorrect classifications, respectively. The size and color intensity of the circle represent the average percentage of assignments into each class based on 100 model-fitting iterations. For species abbreviations see Table S2.

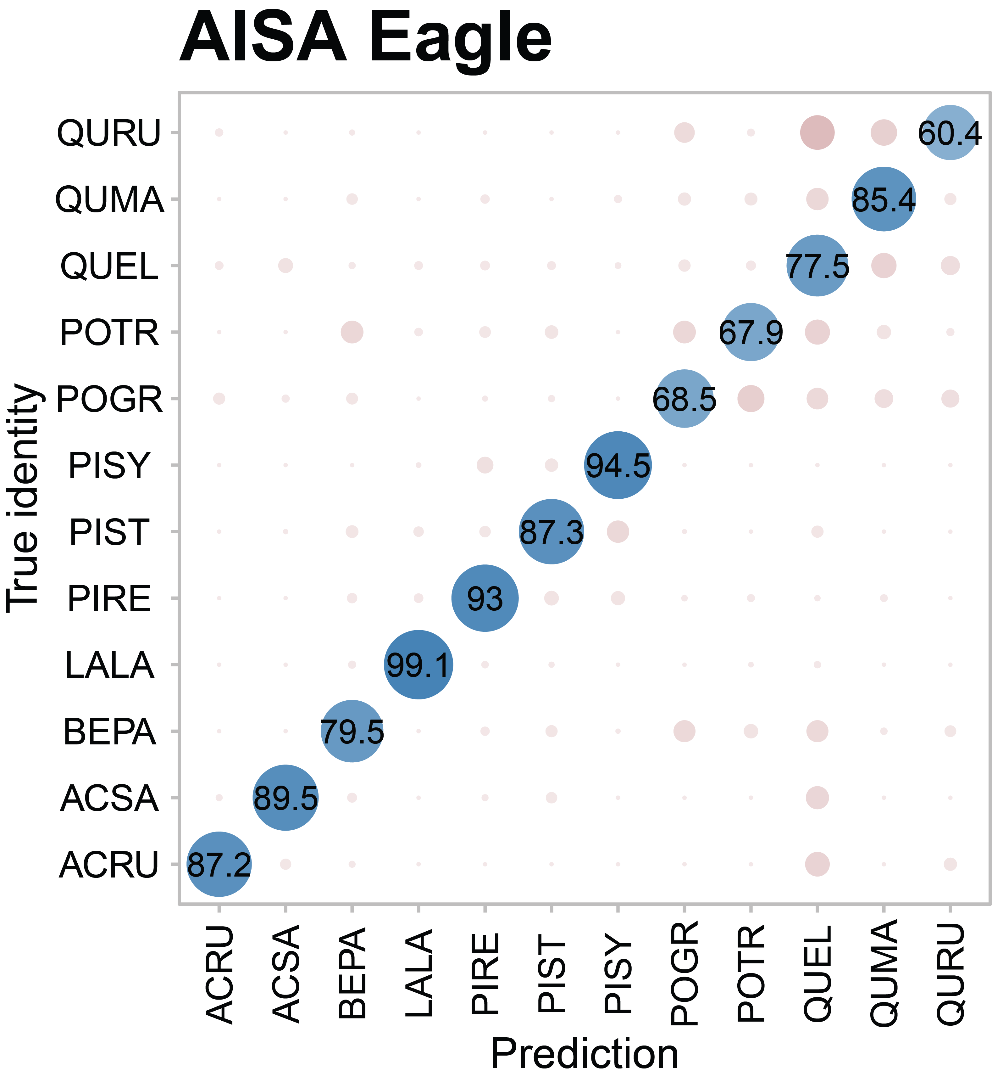

| **AISA Eagle** | | |
| --- | --- | --- |
| **Parameter** | **Mean** | **SD** |
| *Accuracy* | 0.82 | 0.0119 |
| *Kappa* | 0.8 | 0.0133 |
| *AccuracyLower* | 0.79 | 0.0126 |
| *AccuracyUpper* | 0.85 | 0.0111 |
| *AccuracyNull* | 0.18 | <0.0001 |
| *AccuracyPValue* | <0.0001 | <0.0001 |

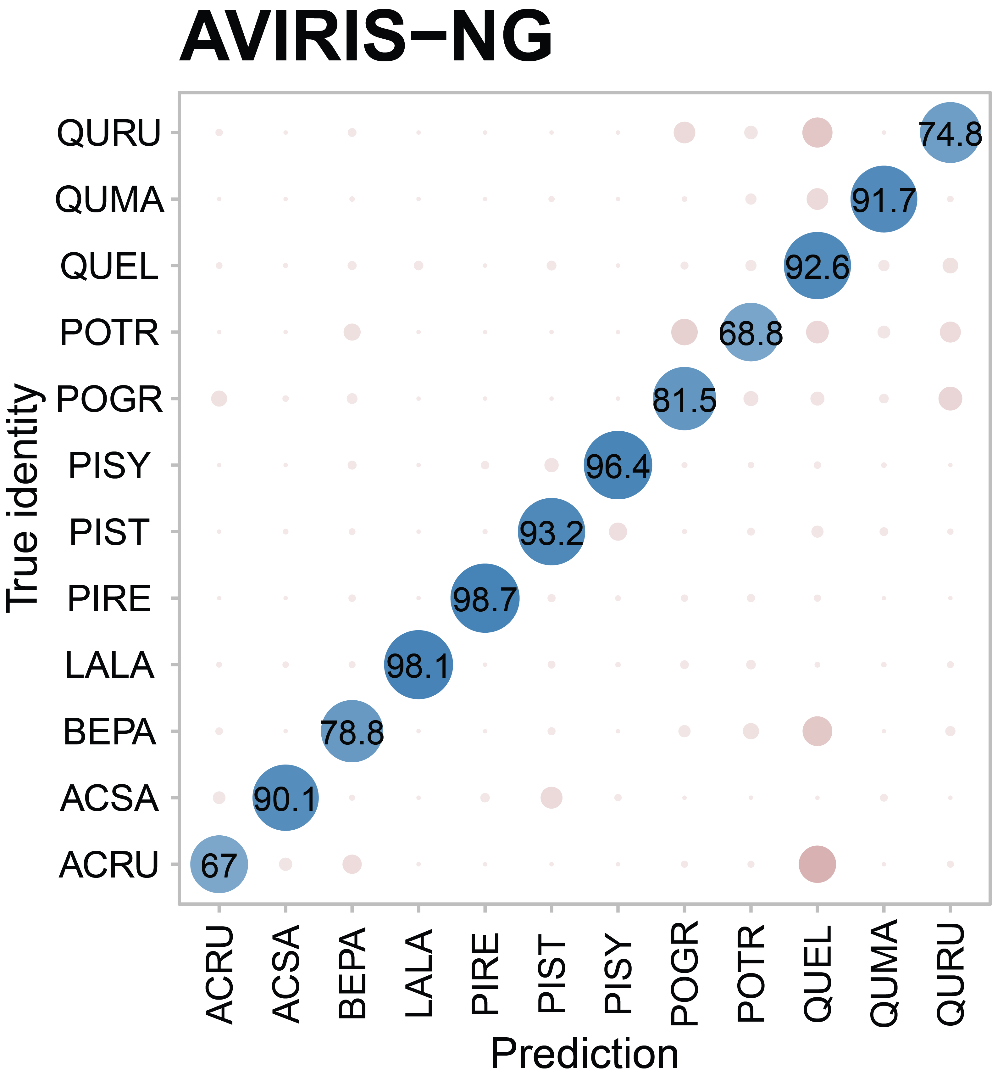

| **AVIRIS-NG** | | |
| --- | --- | --- |
| **Parameter** | **Mean** | **SD** |
| *Accuracy* | 0.9 | 0.0134 |
| *Kappa* | 0.88 | 0.0151 |
| *AccuracyLower* | 0.87 | 0.015 |
| *AccuracyUpper* | 0.92 | 0.0117 |
| *AccuracyNull* | 0.18 | <0.0001 |
| *AccuracyPValue* | <0.0001 | <0.0001 |

**Table S3.** Performance and number of pixels included in each PLSDA.

| PLSDA | # training samples | # testing samples | Producer accuracy (True positive rate) | Omission error (False negative rate) | User accuracy (True negative rate) | Commission error (False positive rate) | Sensitivity | Specificity | Overall accuracy |
| --- | --- | --- | --- | --- | --- | --- | --- | --- | --- |
| *AISA Eagle Direct Diseased vs Other* | 2063 | 688 | 0.19 | 0.81 | 0.99 | 0.01 | 0.19 | 0.99 | 0.59 |
| *AVIRIS-NG Direct Diseased vs Other* | 1324 | 441 | 0.40 | 0.60 | 0.98 | 0.02 | 0.40 | 0.98 | 0.69 |
| *AISA Eagle Phylogenetic Diseased vs Other* | 2063 | 688 | 0.55 | 0.45 | 0.96 | 0.04 | 0.55 | 0.96 | 0.76 |
| *AVIRIS-NG Phylogenetic Diseased vs Other* | 1324 | 441 | 0.71 | 0.29 | 0.97 | 0.03 | 0.71 | 0.97 | 0.84 |
| *AISA Eagle Quercus vs Other Step* | 1547 | 516 | 0.90 | 0.10 | 0.91 | 0.09 | 0.90 | 0.91 | 0.91 |
| *AVIRIS-NG Quercus vs Other Step* | 993 | 331 | 0.99 | 0.01 | 0.95 | 0.05 | 0.99 | 0.95 | 0.97 |
| *AISA Eagle Red vs White Step* | 519 | 173 | 0.91 | 0.09 | 0.85 | 0.15 | 0.91 | 0.85 | 0.88 |
| *AVIRIS-NG Red vs White Step* | 303 | 101 | 0.99 | 0.01 | 0.95 | 0.05 | 0.99 | 0.95 | 0.97 |
| *AISA Eagle Diseased vs Healthy Step* | 345 | 115 | 0.78 | 0.22 | 0.89 | 0.11 | 0.78 | 0.89 | 0.84 |
| *AVIRIS-NG Diseased vs Healthy Step* | 218 | 73 | 0.87 | 0.13 | 0.95 | 0.05 | 0.87 | 0.95 | 0.91 |

**Appendix S3** Performance of oak identification PLS-DA models within the phylogenetic approach for AISA Eagle and AVIRIS-NG VSWIR datasets. Blue and red circles represent correct and incorrect classifications, respectively. The size and color intensity of the circle represent the average percentage of classifications into each group based on 10,000 model-fitting iterations, one standard deviation is shown in parentheses. K values represent the number of components. Models were validated against an independent set of oaks.

**Testing**

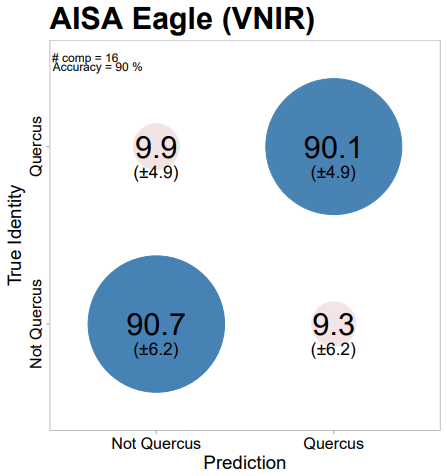

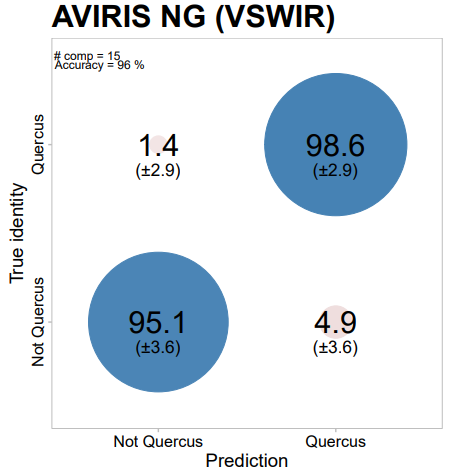

| **AISA Eagle** | | |
| --- | --- | --- |
| **Parameter** | **Mean** | **SD** |
| Accuracy | 0.9 | 0.01 |
| Kappa | 0.79 | 0.0216 |
| AccuracyLower | 0.88 | 0.011 |
| AccuracyUpper | 0.93 | 0.0089 |
| AccuracyNull | 0.67 | <0.0001 |
| AccuracyPValue | <0.0001 | <0.0001 |
| McnemarPValue | 0.11 | 0.2035 |

| **AVIRIS-NG** | | |
| --- | --- | --- |
| **Parameter** | **Mean** | **SD** |
| Parameter | Mean | SD |
| Accuracy | 0.96 | 0.0024 |
| Kappa | 0.91 | 0.0055 |
| AccuracyLower | 0.95 | 0.0027 |
| AccuracyUpper | 0.97 | 0.0021 |
| AccuracyNull | 0.7 | <0.0001 |
| AccuracyPValue | <0.0001 | 0.0009 |

**Validation**

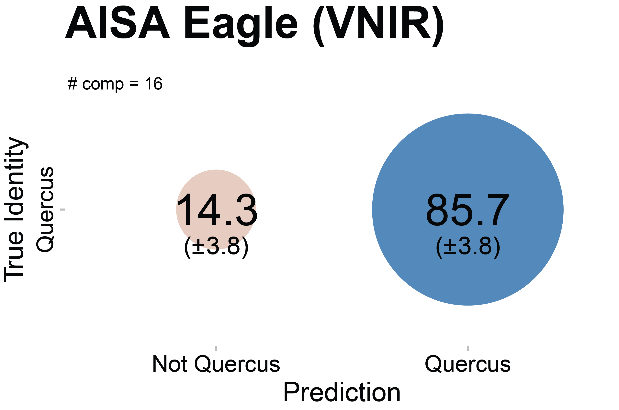

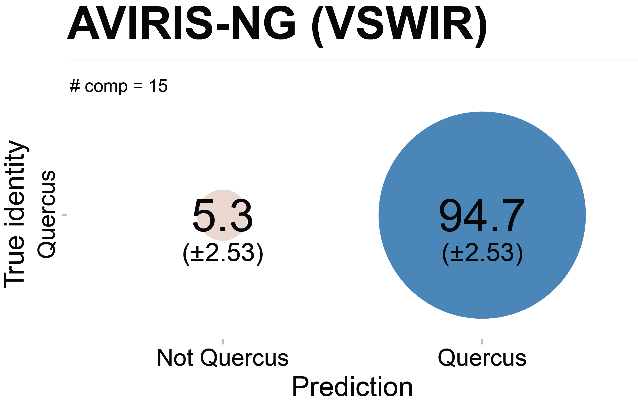

**Appendix S4** Performance of the red oak identification PLS-DA models within the phylogenetic approach for AISA Eagle and AVIRIS-NG VSWIR datasets. Blue and red circles represent correct and incorrect classifications, respectively. The size and color intensity of the circle represent the average percentage of classifications into each group based on 10,000 model-fitting iterations, one standard deviation is shown in parentheses. K values represent the number of components. Models were validated against an independent set of red and white oaks.

**Testing**

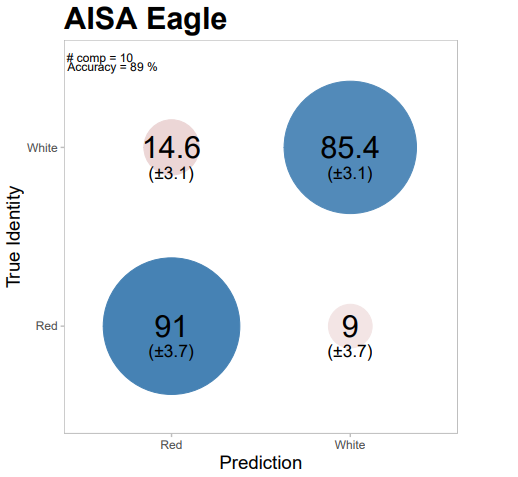

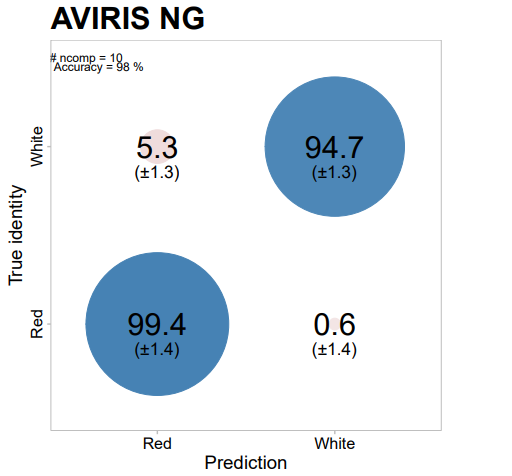

| **AVIRIS-NG** | | |
| --- | --- | --- |
| **Parameter** | **Mean** | **SD** |
| Accuracy | 0.98 | 0.0031 |
| Kappa | 0.95 | 0.0078 |
| AccuracyLower | 0.96 | 0.004 |
| AccuracyUpper | 0.99 | 0.0021 |
| AccuracyNull | 0.72 | <0.0001 |
| AccuracyPValue | <0.0001 | <0.0001 |
| McnemarPValue | 0.25 | 0.2514 |

| **AISA Eagle** | | |
| --- | --- | --- |
| **Parameter** | **Mean** | **SD** |
| Accuracy | 0.89 | 0.0191 |
| Kappa | 0.76 | 0.042 |
| AccuracyLower | 0.84 | 0.0219 |
| AccuracyUpper | 0.93 | 0.0158 |
| AccuracyNull | 0.67 | <0.0001 |
| AccuracyPValue | <0.0001 | <0.0001 |
| McnemarPValue | 0.55 | 0.3286 |

**Validation**

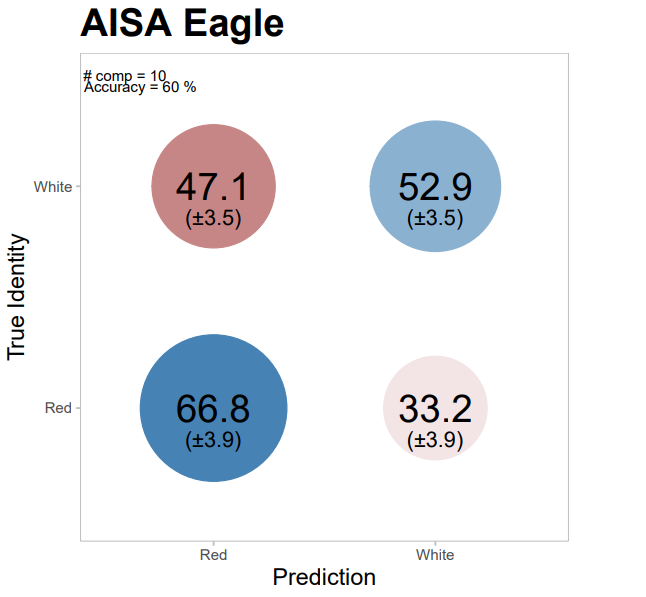

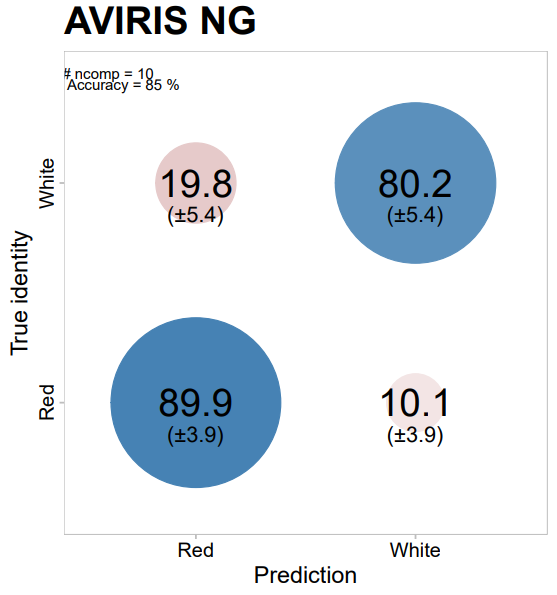

**Appendix S5** Performance of the diseased red oak PLS-DA models within the phylogenetic approach for the AISA Eagle and AVIRIS-NG VSWIR datasets. Blue and red circles represent correct and incorrect classifications, respectively. The size and color intensity of the circle represent the average percentage of classifications into each group based on 10,000 model-fitting iterations, one standard deviation is shown in parentheses. K values represent the number of components.

**Testing**

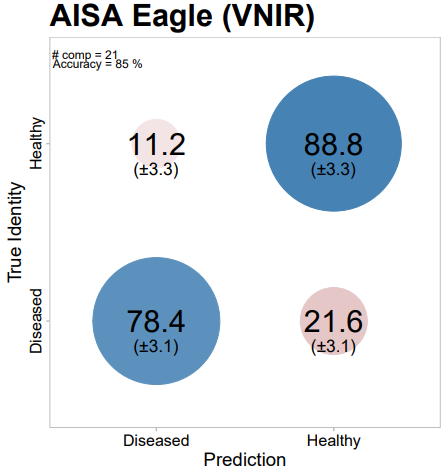

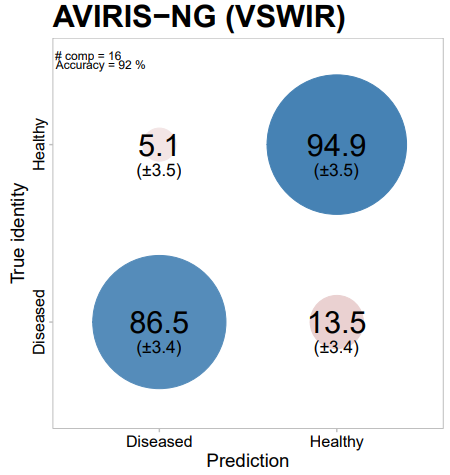

| **AVIRIS-NG** | | |
| --- | --- | --- |
| **Parameter** | **Mean** | **SD** |
| Accuracy | 0.92 | 0.0115 |
| Kappa | 0.82 | 0.0251 |
| AccuracyLower | 0.88 | 0.0132 |
| AccuracyUpper | 0.94 | 0.0095 |
| AccuracyNull | 0.64 | <0.0001 |
| AccuracyPValue | <0.0001 | <0.0001 |
| McnemarPValue | 0.42 | 0.3392 |

| **AISA Eagle** | | |
| --- | --- | --- |
| **Parameter** | **Mean** | **SD** |
| Accuracy | 0.85 | 0.0266 |
| Kappa | 0.68 | 0.057 |
| AccuracyLower | 0.78 | 0.0301 |
| AccuracyUpper | 0.9 | 0.022 |
| AccuracyNull | 0.62 | <0.0001 |
| AccuracyPValue | <0.0001 | <0.0001 |
| McnemarPValue | 0.57 | 0.3258 |

**Table S4.** Performance of PLS-DAs when applied to data obtained with a sensor other than the one used for training.

| PLSDA | Producer accuracy (True positive rate) | Omission error (False negative rate) | User accuracy (True negative rate) | Commission error (False positive rate) | Sensitivity | Specificity | Overall accuracy |
| --- | --- | --- | --- | --- | --- | --- | --- |
| *AISA Eagle Quercus vs Other Step on AVIRIS-NG VNIR data* | 0.64 | 0.36 | 0.36 | 0.64 | 0.64 | 0.36 | 0.50 |
| *AVIRIS-NG VNIR Quercus vs Other Step on AISA Eagle data* | 0.16 | 0.84 | 0.76 | 0.24 | 0.16 | 0.76 | 0.46 |
| *AISA Eagle Red vs White Step on AVIRIS-NG VNIR data* | 0.35 | 0.65 | 0.64 | 0.36 | 0.35 | 0.64 | 0.50 |
| *AVIRIS-NG VNIR Red vs White Step on AISA Eagle data* | 0.74 | 0.26 | 0.26 | 0.74 | 0.74 | 0.26 | 0.50 |
| *AISA Eagle Diseased vs Healthy Step on AVIRIS-NG VNIR data* | 0.21 | 0.79 | 0.79 | 0.21 | 0.21 | 0.79 | 0.50 |
| *AVIRIS-NG VNIR Diseased vs Healthy Step on AISA Eagle data* | 0.58 | 0.42 | 0.41 | 0.59 | 0.58 | 0.41 | 0.50 |

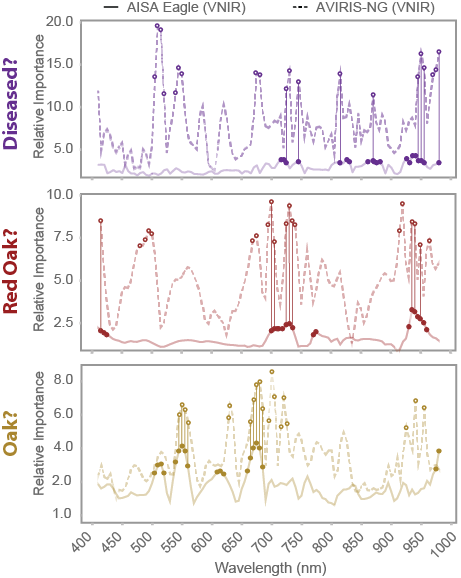

**Figure S3.** AISA Eagle (solid line, closed circles) and AVIRIS-NG (dashed line, open circles) sensors share wavelengths that are important to distinguish oaks, red oaks, and diseased red oaks. We trained PLS-DA models for each sensor using the same range of wavelengths (VNIR, 410-980 nm) to identify important wavelengths that are common (circles connected with vertical lines) in both sensor platforms. The oak models share nine wavelengths located around 550 nm and 675 nm. The red oak models share nine wavelengths located at 410 nm, between 700-750 nm, and around 950 nm. The diseased red oak models share eight wavelengths between 770-750 nm, 800-825 nm, 850-900 nm, around 950 nm, and at 980 nm.

**Table S4.** Differences in oak wilt detectability between late July and late August. F-values and significances for each index correspond to pairwise comparisons between healthy and diseased red oak canopy pixels.

|  | **AISA Eagle-July** | | **AVIRIS-NG-August** | |
| --- | --- | --- | --- | --- |
| **Index** | **F-value** | **Significance** | **F-value** | **Significance** |
| *CMS* | 123.45 | *** | 23.31 | *** |
| *PRIn* | 61.5 | *** | 1.62 |  |
| *NPCI* | 7.1 | ** | 3.66 | . |
| *SIPI* | 89.86 | *** | 22.18 | *** |
| *WBI* | 30.24 | *** | 35.08 | *** |
| *SR* | 67.48 | *** | 38.44 | *** |
| *PRI570* | 56.97 | *** | 0.61 |  |
| *RDVI* | 119.88 | *** | 38.49 | *** |
| *CI* | 121.81 | *** | 28.31 | *** |
| *NPQI* | 0.14 |  | 0.01 |  |
| *PRIxCI* | 46.39 | *** | 0.01 |  |
| *PRIm1* | 27.51 | *** | 0 |  |
| *CRI700* | 68.22 | *** | 4.09 | * |
| *PRIm4* | 5.06 | * | 8.87 | ** |
| *DCabCxc* | 20.79 | *** | 0.99 |  |
| *VOG2* | 89.8 | *** | 21.95 | *** |
| *TCARI/OSAVI* | 72.15 | *** | 11.48 | *** |
| *NDWI* | - | - | 51.36 | *** |
| *WBI SWIR* | - | - | 41.94 | *** |
| *SR_SIF_* | 119.98 | *** | 24.03 | *** |
| *CCI* | 17.09 | *** | 17.12 | *** |

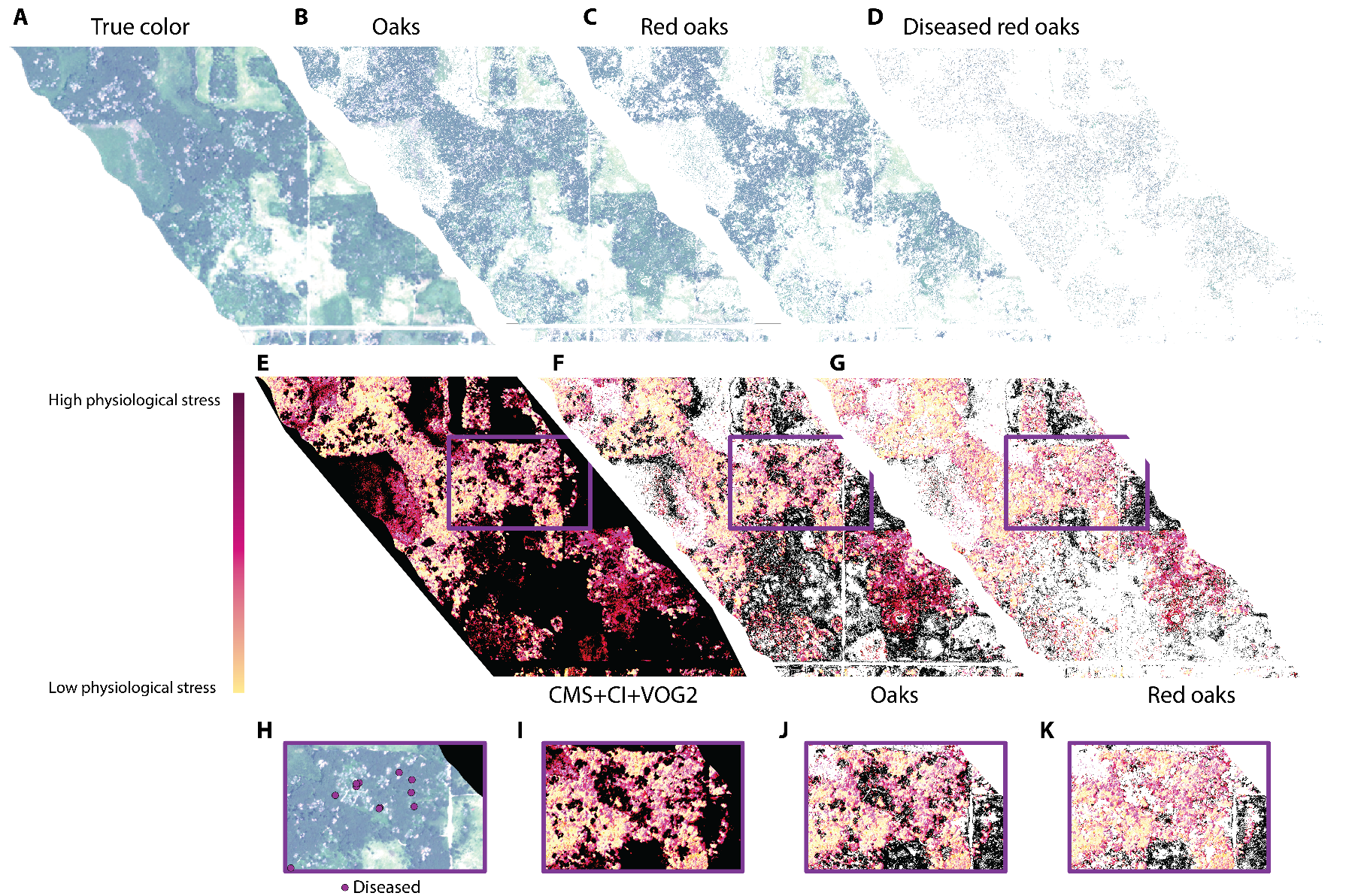

**Figure S4.** Mapping of the AISA Eagle PLS-DA models. The Oak discrimination model was applied to the full scenery and the resulting map was used to mask out pixels with a probability of being an oak lower than 0.8. The red oak model was applied to the remaining pixels and pixels with a probability of being a red oak lower than 0.8 were masked out. Finally, the diseased model was applied to the remaining pixels and pixels with a probability of being a diseased red oak lower than 0.8 were masked out. Spectral indices are then used to evaluate areas with pixels classified as potential diseased red oaks by PLS-DA models.

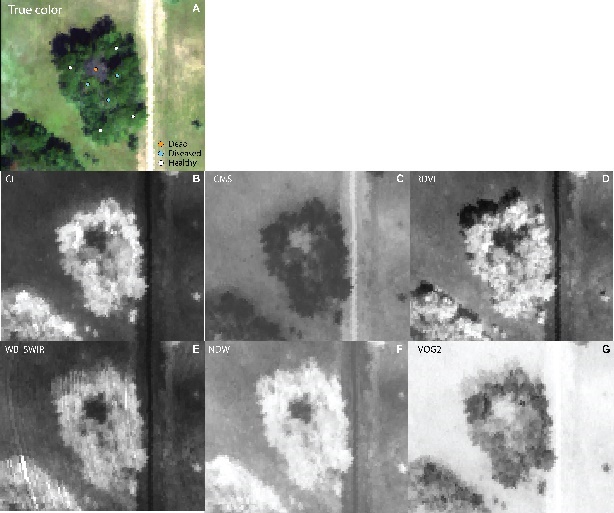
**Figure S5.** A typical oak wilt pocket observed through different spectral indices using the 2016 AVIRIS-NG data. A tree killed by oak wilt during 2015 can be observed at the center of the oak wilt pocket. Three diseased trees stand next to it. Diseased trees remain undetectable in true color images (red as 640 nm, green as 550 nm, and blue as 470 nm) (A). However, they become apparent through spectral indices associated with photosynthetic function (B-D), such as CI, CMS, or RDVI; water status (E-F), such as NDWI or WBI SWIR; or indices sensitive to both, such as VOG2 (G). See Table S1 for index abbreviations.

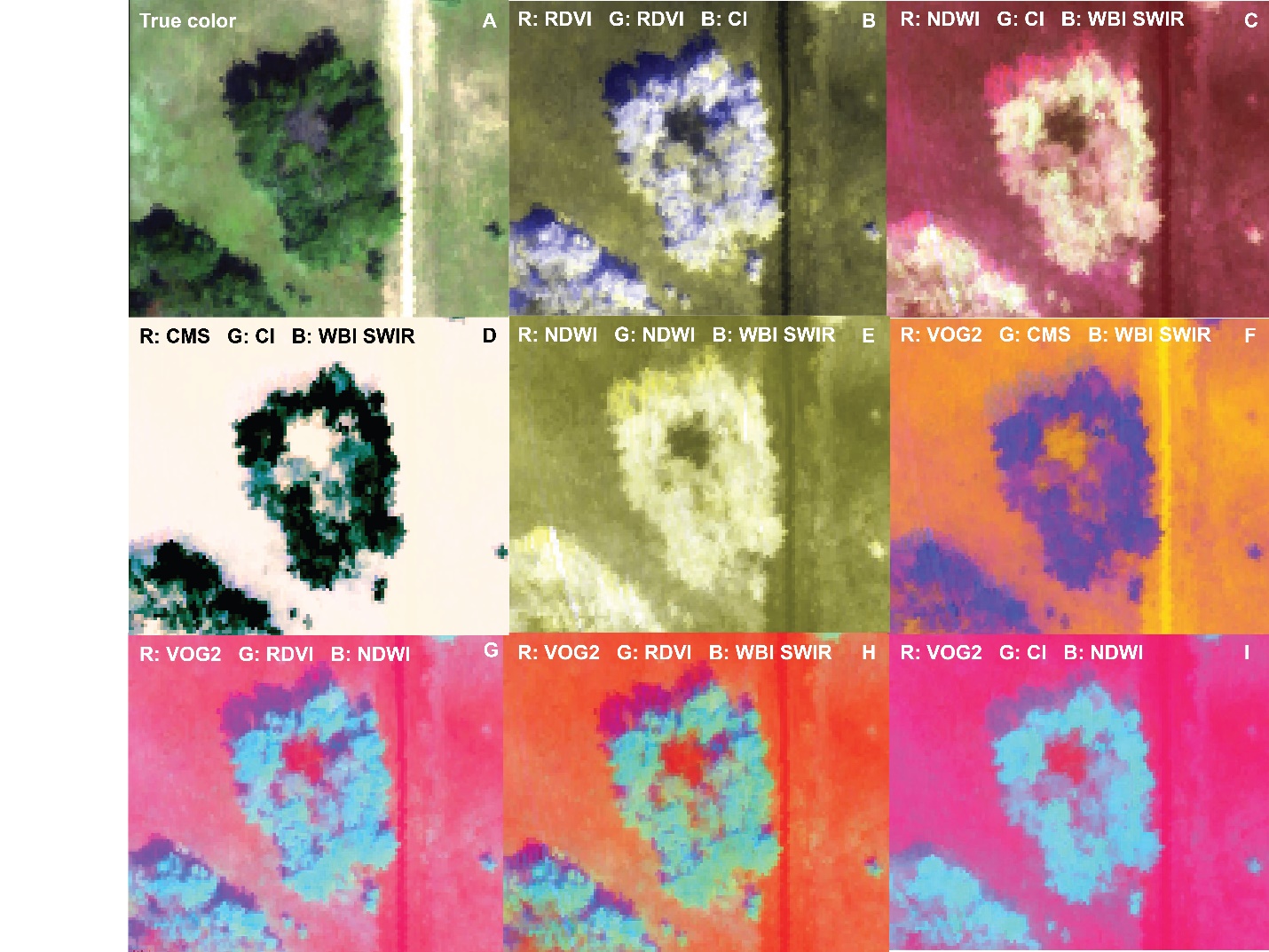

**Figure S6.** A typical oak wilt pocket observed through different combinations of spectral indices using the 2016 AVIRIS-NG data. A tree killed by oak wilt during 2015 can be observed at the center of the oak wilt pocket in true color (red as 640 nm, green as 550 nm, and blue as 470 nm) (A). Three diseased trees stand next to it that cannot be detected with true color images. However, they become apparent through spectral indices associated with photosynthetic function and water status placed (B-I) on the red (R), green (G), and blue (B) channels. See Table S1 for index abbreviations.
